# Suppression of membranous LRP5 recycling, WNT/β-catenin signaling, and colorectal tumorigenesis by 15-LOX-1 peroxidation of PI3P_linoleic acid

**DOI:** 10.1101/747592

**Authors:** Fuyao Liu, Xiangsheng Zuo, Yi Liu, Yasunori Deguchi, Micheline J. Moussalli, Weidong Chen, Peiying Yang, Bo Wei, Lin Tan, Philip L. Lorenzi, Shen Gao, Jonathan C. Jaoude, Amir Mehdizadeh, Lovie Ann Valentin, Daoyan Wei, Imad Shureiqi

**Author notes:** Correspondence (I.S.). Department of Gastrointestinal Medical Oncology, the University of Texas MD Anderson Cancer Center, Houston, TX 77030. These authors contributed equally to this manuscript (Fuyao Liu, Xiangsheng Zuo).

## Abstract

Aberrant Wnt/β-catenin activation is a major driver of colorectal cancer (CRC), which is typically initiated by *APC* mutations. Additional modifiable factors beyond *APC* mutations have been recognized to be important for further potentiation of aberrant β-catenin activation to promote colorectal tumorigenesis. These factors have yet to be clearly identified. Western-type diets are increasingly enriched in linoleic acid (LA). LA-enriched diet however promotes chemically-induced colorectal tumorigenesis in rodent models. Furthermore, the main metabolizing enzyme of LA, 15-lipoxygenase-1 (15-LOX-1), is transcriptionally silenced in CRC. Whether LA and 15-LOX-1 affect Wnt/β-catenin signaling to modulate colorectal tumorigenesis is poorly understood. Herein, we report that high dietary LA promoted colorectal tumorigenesis in mice with intestinally targeted *APC* mutation (*Apc^Δ580^*) by upregulating a Wnt receptor, LRP5 expression, and β-catenin activation. 15-LOX-1 transgenic expression in intestinal epithelial cells suppressed LRP5 expression, β-catenin activation and subsequently CRC in these mice. In particular, 15-LOX-1 peroxidation of LA in phosphatidylinositol-3-phosphates (PI3P_LA) into PI3P_13-HODE decreased PI3P binding to SNX17and LRP5, which inhibited LRP5 recycling from endosomes to the plasma membrane, thereby leading to an increase of LRP5 lysosomal degradation. Our findings demonstrate for the first time that 15-LOX-1 metabolism of LA in PI3P to regulate LRP5 membrane abundance is a modifiable factor of Wnt/β-catenin aberrant signaling that could be potentially therapeutically targeted to suppress colorectal tumorigenesis and progression.

## Introduction

Colorectal cancer (CRC) is the third leading cause of cancer deaths in the United States (Siegel et al., 2014). *APC* mutations, which occur early in most CRCs (Losi et al., 2005), increase β-catenin activation, driving the initiation of colorectal carcinogenesis (Fearon and Vogelstein, 1990, Clevers and Nusse, 2012). β-catenin activation varies significantly even within the same CRC that has the same *Apc* mutation (Vermeulen et al., 2010, Brabletz et al., 1998), and this variability has been attributed to modifiable factors that can strongly impact colorectal tumorigenesis and progression via β-catenin hyperactivation (He et al., 2005, Suzuki et al., 2004, Vermeulen et al., 2010). Unlike *APC* mutations, which are not modifiable by current therapeutic approaches, the identification of these modifiable factors could open important therapeutic targeting opportunities.

Linoleic acid (LA) is the most commonly consumed n-6 polyunsaturated fatty acid in Western diets (Adam et al., 2008, Shureiqi et al., 2010) and promotes CRC in chemically-induced carcinogenesis models in rodents (Deschner et al., 1990, Lipkin et al., 1999). Nevertheless, human diets, especially in the United States, have been increasingly enriched with LA during the past six decades in response to the notion that LA decreases the risk of coronary artery disease (Farvid et al., 2014, Blasbalg et al., 2011). This notion has recently been challenged by new findings showing that substitution of dietary saturated fats with LA increased cardiovascular disease risk (Ramsden et al., 2013). While this debate continues, the effect of dietary LA on CRC risk remains poorly defined. Results of dietary LA studies in human CRC remain inconclusive (Jandacek, 2017) possibly because they depend on participants’ recall of dietary intake. In preclinical models, however, where dietary factors can be strictly controlled, LA promoted azoxymethane (AOM)–induced CRC in rodents (Lipkin et al., 1999, Deschner et al., 1990). Oxidative metabolism of polyunsaturated fatty acids such as LA influences their ability to promote AOM-induced CRC in rodents (Bull et al., 1989). LA’s oxidative metabolism occurs mainly via a pathway regulated by 15-lipoxygenase-1 (15-LOX-1) to generate 13-S-hydroxyoctadecadienoic acid [13(S)-HODE]. 15-LOX-1 is downregulated in more than 60% of advanced colorectal adenomas and 100% of human invasive CRCs (Yuri et al., 2007, Shureiqi et al., 1999); and re-expression of 15-LOX-1 expression suppresses CRC in various preclinical models (Shureiqi et al., 2003, Nixon et al., 2004, Shureiqi et al., 2005, Wu et al., 2008, Zuo et al., 2012, Mao et al., 2015).

Whether this alteration in LA oxidative metabolism via 15-LOX-1 impacts aberrant Wnt/β-catenin signaling in CRC is poorly defined because the few available findings are derived from studies of 12/15-LOX in non-cancer mouse models, which have produced conflicting results suggesting that 12/15-LOX may either inhibit (Almeida et al., 2009) or stimulate (Kinder et al., 2010) β-catenin transcriptional activity. Experimental modeling of 15-LOX-1 via its murine homologue, 12/15-LOX, is suboptimal because 12/15-LOX is a hybrid enzyme of 15- and 12-LOX, whose products have opposing biological effects on important physiologic and pathologic processes, especially in the case of tumorigenesis (Liu et al., 1995, Muller et al., 2002). Thus, important questions remain: do LA and its oxidative metabolism by 15-LOX-1 impact Wnt/β-catenin signaling and subsequently CRC? If so, by what mechanisms? These questions are important because increasing dietary LA intake as advocated to reduce the risk of coronary artery disease (Farvid et al., 2014) could inadvertently enhance the risk of CRC, especially when 15-LOX-1 is downregulated, as commonly occurs in CRC (Yuri et al., 2007, Shureiqi et al., 1999). The necessity of addressing those questions is underlined by the recent findings showing that the oxidative metabolism of LA via cytochrome P450 monooxygenases to produce 12,13-epoxyoctadecenoic acid, which likely occurs in the absence of 15-LOX-1, contributes to CRC promotion (Wang et al., 2019).

In this study, we addressed these questions in complementary human and mouse experimental models and found that: 1) high dietary LA promotes CRC by upregulating the expression of a Wnt receptor, low-density lipoprotein (LDL) receptor–related protein 5 (LRP5) to increase aberrant β-catenin activation in intestinal epithelial cells (IECs); and 2) 15-LOX-1 re-expression in IECs suppresses CRC by peroxidation of LA in phosphatidylinositol-3-phosphates (PI3P_LA) into PI3P_13-HODE, which decreases PI3P binding to sorting nexin 17 (SNX17) and LRP5 and subsequently inhibits LRP5 recycling from endosomes to the plasma membrane, thus leading to an increase of LRP5 lysosomal degradation. Thus our findings uncover a regulatory mechanism of LRP5 membrane abundance as a modifiable factor of Wnt/β-catenin aberrant signaling that modulates CRC tumorigenesis and progression.

## Results

### 15-LOX-1 suppresses LA promotion of *Apc* mutation–induced CRC and aberrant β-catenin activation in mice

We first examined the effects of 15-LOX-1 transgenic expression (Supplementary Figure 1A) on dietary LA promotion of AOM–induced CRC by feeding *15-LOX-1*-Gut mice (with 15-LOX-1 transgenic overexpression in IECs (Zuo et al., 2012)) and their wild-type (WT) littermates with standardized iso-caloric diets in which corn oil (containing 50%-55% LA) comprised either 5% of diet weight, considered low, or 20% of diet weight, which simulates the high LA content of Western-type diets (Deschner et al., 1990, Yang et al., 1996). In WT mice, AOM induced significantly more colorectal tumors per mouse in those fed 20% corn oil (mean ± standard error: 7.11 ± 1.37) than in those fed 5% corn oil (3.33 ± 1.56) (Supplementary Figures 1B and C). In contrast, in *15-LOX-1*-Gut mice, the numbers of tumors per mouse were similar between those fed 20% corn oil (2.5 ± 0.82) and those fed 5% corn oil (2.8 ± 1.87). Moreover the *15-LOX-1*-Gut mice had significantly fewer tumors per mouse than did the corresponding WT littermates fed the same diets.

To ensure that LA promotion of CRC and its modulation by 15-LOX-1 are not limited to the AOM carcinogenesis model, we examined the effects of 15-LOX-1 and the same low and high corn-oil diets on *Apc* mutation–driven CRC using *Apc^Δ580^* mice without and with transgenic 15-LOX-1 expression in IECs (*Apc ^Δ580^–15-LOX-1* mice) (Supplementary Figures 1D-F). In *Apc^Δ580^*mice, increasing the corn-oil content from 5% to 20% significantly increased body weight (Supplementary Figure 1G) and tumor burden at age 14 weeks (Figures 1A and B). In contrast, *Apc^Δ580^–15-LOX-1* mice had a lower tumor burden than did the *Apc^Δ580^* mice when fed either 5% or 20% corn oil, especially for large tumors (diameter>3 mm) (Figure 1C), despite a body weight increase in those fed 20% corn oil compared with those fed 5% corn oil (Supplementary Figure 1G). In a longitudinal follow-up survival experiment, *Apc^Δ580^–15-LOX-1* mice lived significantly longer than did the corresponding *Apc^Δ580^* littermates fed the same diets; at 47 weeks, none of the *Apc^Δ580^* mice were alive, while 53% of the *Apc^Δ580^–15-LOX-1* mice fed 5% corn oil and 25% of the *Apc^Δ580^*–15-LOX-1 mice fed 20% corn oil were still alive (Figure 1D). To determine whether 15-LOX-1 impacts β-catenin and CRCs even in the absence of dietary LA variation, we fed *Apc^Δ580^–15-LOX-1* mice and their *Apc^Δ580^* littermates the same 7% corn-oil diet. *Apc^Δ580^–15-LOX-1* mice had markedly higher 13-HODE levels but lower active β-catenin levels and colorectal tumor multiplicity, especially for large tumors (diameter>3 mm) (Supplementary Figures 1H-K), than did *Apc*^Δ580^ littermates at age 25 weeks.

**Figure 1.**
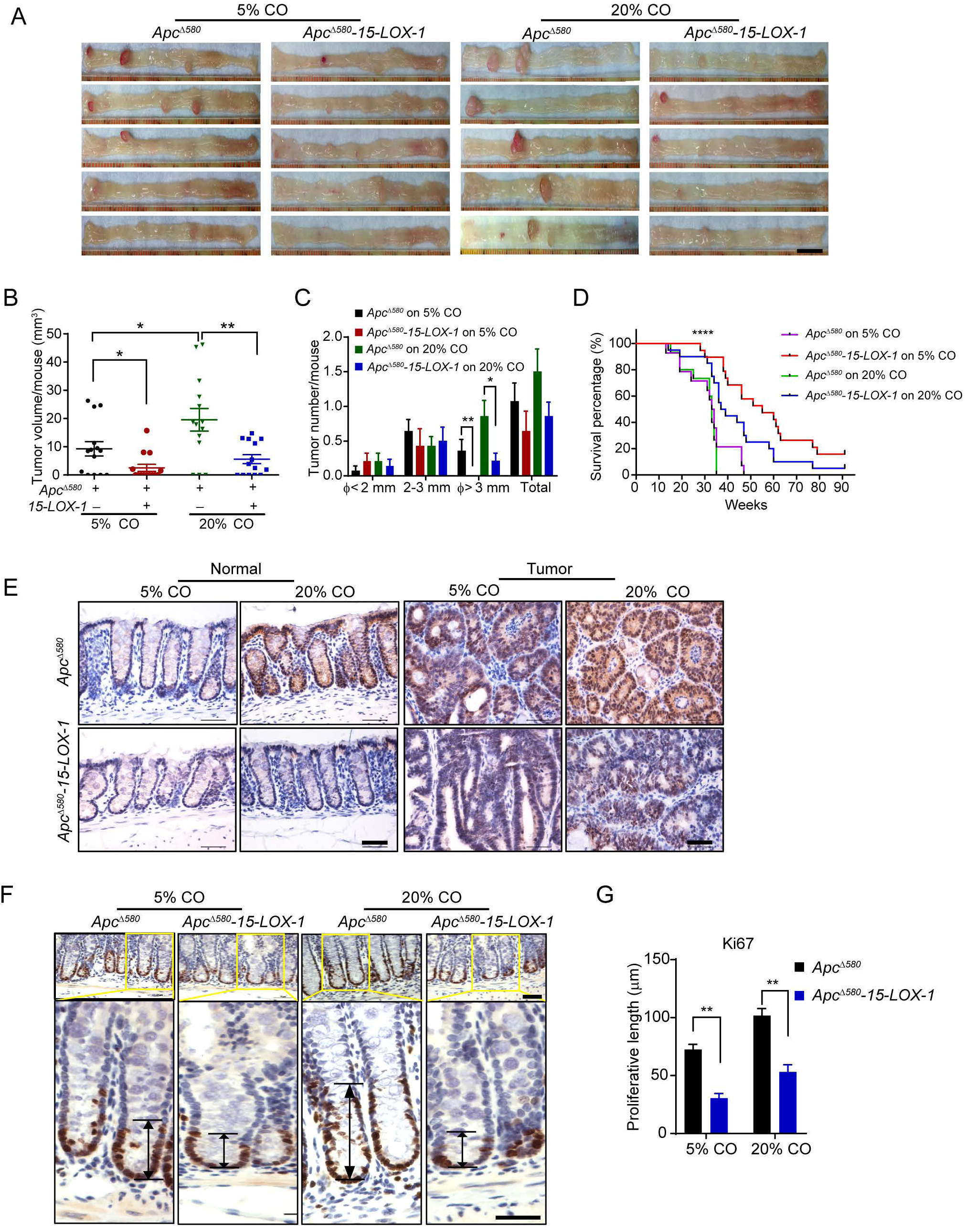
15-LOX-1 suppresses linoleic acid (LA) promotion of *Apc* mutation–induced colorectal cancer (CRC) and aberrant β-catenin activation in mice. **(A-C)** *Apc*^Δ580^ and *Apc^Δ580^–15-LOX-1* littermates at 4 weeks were fed either 5% or 20% corn oil (CO), killed at age 14 weeks, and examined for tumor formation (n=14 mice per group). Representative colon images **(A),** colonic tumor volumes per mouse **(B),** and colonic tumor numbers and size distributions per mouse **(C)** of the indicated mouse groups. **(D)** Survival curves of *Apc^Δ580^* and *Apc^Δ580^–15-LOX-1* littermates at 4 weeks fed either 5% or 20% corn oil (n=14-20 mice per group). **(E)** Representative immunohistochemistry staining images of active β-catenin expression and localization in normal and tumor colonic tissues from the indicated mouse groups as described in **A-C**. **(F, G)** Representative immunohistochemistry staining images for cell proliferation marker Ki-67 **(F)** and corresponding proliferation zone lengths **(G)** in normal colonic tissues from the indicated mouse groups as described in **A-C**. Scale bars: 1 cm (A) and 100 µm (E and F)

*Apc^Δ580^* mutation induces aberrant β-catenin activation to initiate CRC tumorigenesis in *Apc^Δ580^*mice^32^, we therefore examined whether dietary LA and 15-LOX-1 further modulate β-catenin activation and affect CRC in these mice. β-catenin levels in both normal-appearing and tumor mucosa were increased by high dietary corn oil but decreased by 15-LOX-1 transgenic expression in IECs (Figure 1E). *Apc* mutations have been reported to expand the colonic crypt proliferative zones as an important mechanism to drive CRC (Barthold and Beck, 1980, Boman and Fields, 2013); in our study, colonic crypt proliferative zones as measured by Ki-67 immunohistochemical analysis were increased by high dietary corn oil but decreased by transgenic 15-LOX-1 expression in IECs (Figures 1F and G). Together, these data implicate that 15*–*LOX-1-mediated metabolism of LA modulates β-catenin*–*driven CRC tumorigenesis and progression.

### 15-LOX-1 suppresses LA upregulation of LRP5 and active β-catenin in IECs of *Apc^Δ580^* mice and in human CRC cells

LRPs have dual receptor roles: 1) LA-enriched LDL endocytosis and subsequent degradation (Spiteller and Spiteller, 2000); and 2) transduction of Wnt signals to activate β-catenin (Zhu et al., 2003, Tortelote et al., 2017). In their LDL-related role, LRPs recruit 12/15-LOX to the cell membrane to oxidize LDL (Han et al., 2015). We therefore examined the effects of LA and 15-LOX-1 on these dual roles of LRPs using LRP5 as a representative member.

The LA-enriched high corn oil diet upregulated, while transgenic 15-LOX-1 expression in IECs downregulated, LRP5 and active β-catenin levels in *Apc^Δ580^* mice at age 14 weeks (Figures 2A-C). We also found LRP5 and active β-catenin levels were lower in *Apc^Δ580^*–*15-LOX-1* mice than in *Apc^Δ580^* mice at age 6 to 8 weeks, which precedes the development of CRC in these mice (Supplementary Figure 2A). In *Apc^Δ580^* mice, mRNA levels of β-catenin downstream targets (*Axin2* and *cyclin D1*) were increased by higher LA dietary contents, but decreased by 15-LOX-1 transgenic expression in IECs (Supplementary Figures 2B and C).

**Figure 2.**
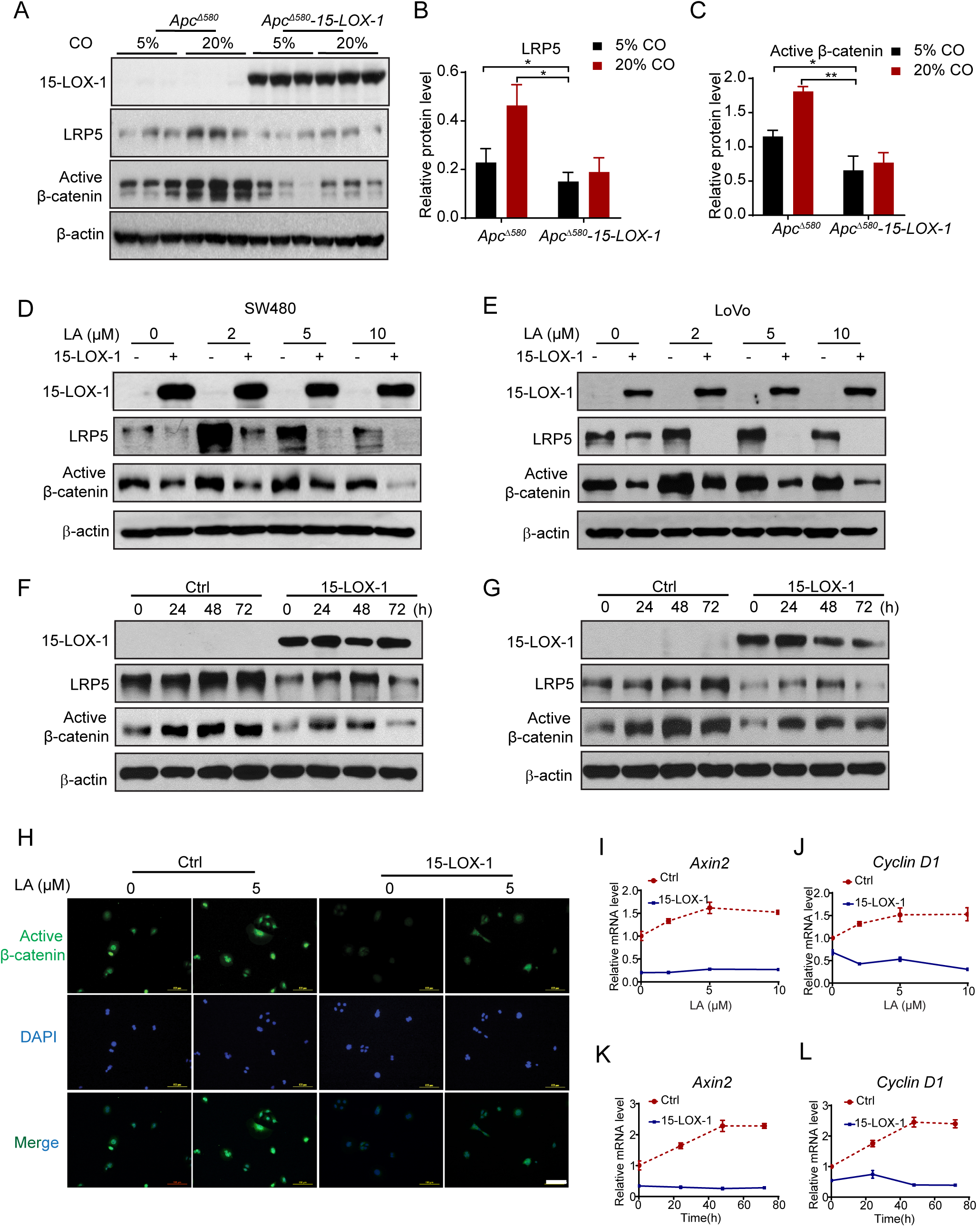
15-LOX-1 suppresses LA upregulation of LRP5 and active β-catenin expression in intestinal epithelial cells (IECs) of *Apc^Δ580^* mice and in human CRC cells. **(A)** Western blot images showing 15-LOX-1, LRP5, and active β-catenin protein expression in IECs from the indicated mouse groups as described in Figures 1A-C. **(B, C)** The band density ratios of LRP5 **(B)** and active β-catenin **(C)** to β-actin corresponding to panel **A**, measured by ImageJ. **(D-G)** Active β-catenin and LRP5 protein levels as measured by Western blot in SW480 and LoVo cells stably transduced with either control (Ctrl) or 15-LOX-1 lentivirus and treated with LA at different indicated concentrations in culture media for 72 hours (**D, E**) or treated with 5 µM LA at different indicated time points (**F, G**). **(H)** Representative immunofluorescence staining images for active β-catenin in SW480 cells treated with 5 µM LA for 48 hours. **(I-L)** *Axin2* and c*yclin D1* mRNA expression in SW480 cells stably transduced with either control (Ctrl) or 15-LOX-1 lentivirus and treated with LA at different concentrations in culture medium supplemented with 5% dialyzed FBS for 48 hours (**I, J**) or treated with 5 µM LA at different time points **(K, L),** measured by qRT-PCR. Scale bars: 100 µm (H).

We next sought to determine whether our mouse findings were applicable to human CRC by re-expressing 15-LOX-1 via lentivirus infection in SW480 and LoVo cells, which, like other CRC cells, lack 15-LOX-1 expression (Moussalli et al., 2011). In the control SW480 and LoVo cells, LA supplementation increased protein levels of LRP5 and active β-catenin, which peaked at 2 and 5 µM LA after 48 to 72 hours (Figures 2D-G). β-catenin nuclear localization (Figure 2H; Supplementary Figure 2D) and *Axin2* and *cyclin D1* mRNA levels increased after LA supplementation in a time- and concentration-dependent manner (Figures 2I-L, and Supplementary Figures 2E-H). All these effects were inhibited by 15-LOX-1 re-expression in SW480 and LoVo cells (Figures 2D-L; Supplementary Figures 2D-H). These observations implicate 15-LOX-1 as a negative modulator of LA-increased LRP5 expression and β-catenin activation.

### 15-LOX-1 inhibits colonic stem cell regeneration in organoids derived from *Apc^Δ580^* mice and human CRC

We assessed the biological significance of 15-LOX-1’s suppression of LRP5 and β-catenin signaling by examining how 15-LOX-1 affects colonic tumor stem cell self-renewal, which is strongly enhanced by aberrant β-catenin activation (Schwitalla et al., 2013). Colonic organoids derived from IECs of *Apc^Δ580^–15-LOX-1* mice had significantly lower expression of LRP5, active β-catenin, Axin2, and cyclin D1 (Figures 3A-C), and fewer primary (P0) and secondary (P1) organoid numbers, especially primitive spheroids, than the ones derived from IECs of Apc^Δ580^ mice did (Figures 3D-F). More importantly, 15-LOX-1 re-expression via lentivirus infection in human CRC organoids derived from patients’ CRC tissues reduced LRP5, active β-catenin, Axin1 and cyclin D1 protein levels (Figures 3G and H), decreased *Axin2* and *cyclin D1* mRNA expression (Figures 3I and J), and repressed their self-renewal, as measured by organoid formation assay (Figures 3K and L).

**Figure 3.**
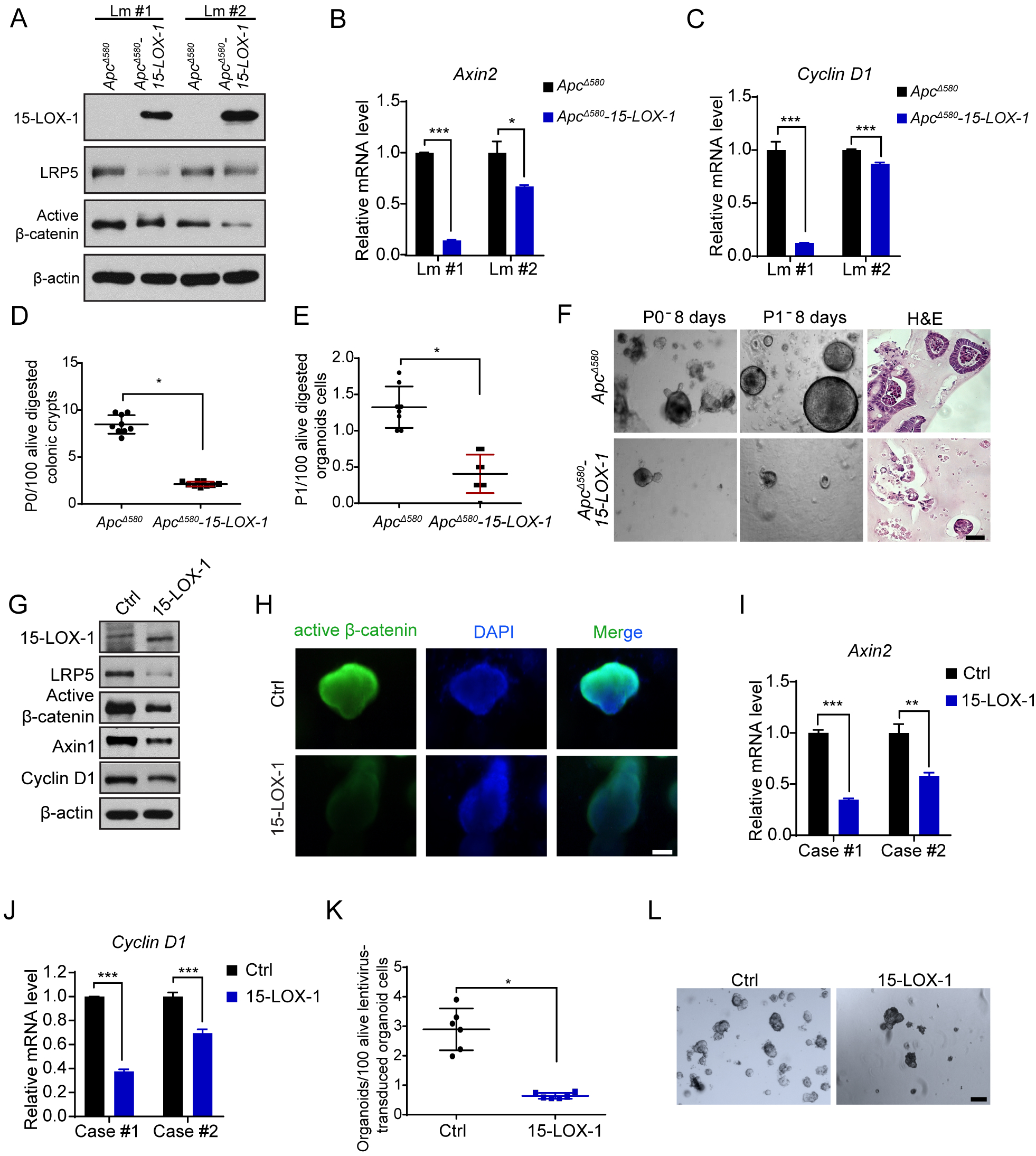
15-LOX-1 inhibits colonic stem cell self-renewal in IECs of *Apc^Δ580^* mice and human CRC cells. **(A-C)** *Apc*^Δ580^ and *Apc*^Δ580^*–15-LOX-1* littermates were killed at 6 weeks, and colonic mucosa tissues were digested and processed for primary organoid cultures for 8 days. LRP5, active β-catenin, and 15-LOX-1 protein expression levels **(A)**, *Axin2* **(B)** and *cyclin D1* **(C)** mRNA expression in the organoid cells from two littermates (Lm) of the indicated mice. **(D-F)** Organoid formation efficacies of primary (P0) **(D)** and secondary (P1) **(E)** organoids cultured for 8 days for the indicated mice. **(F)** Representative bright field light microscopic and hematoxylin and eosin (H&E) staining images (P1) of organoids for the indicated mice. **(G-L)** LRP5, active β-catenin, 15-LOX-1, Axin1 and cyclin D1 protein expression levels, measured by Western blot **(G)**; immunofluorescence staining images of active β-catenin **(H),** and mRNA expression of *Axin2* **(I)** and *cyclin D1* **(J)**, measured by qRT-PCR in human CRC organoid cells transduced with control (Ctrl) or 15-LOX-1 lentivirus. (**K**) Organoid formation efficacies of human CRC–derived organoids transduced with either control or 15-LOX-1 lentivirus. **(L)** Representative bright field light microscopic images for human CRC organoids transduced with control (Ctrl) or 15-LOX-1 lentivirus. Scale bars: 100 µM (F, H, and L).

### 15-LOX-1 downregulates LRP5 expression via 13-HODE

To identify the molecular mechanisms by which 15-LOX-1 downregulates LRP5 expression, we first examined the oxidative lipid profiles of IECs via liquid chromatography–tandem mass spectrometry (LC-MS/MS) to determine the relationship between 15-LOX-1 enzymatic products and their suppression of CRC in *Apc^Δ580^* mice. In *Apc^Δ580^–15-LOX-1* mice, the dominant oxidative lipid products were 13-HODE and 15-Hydroxyeicosatetraenoic (HETE) acid (Figure 4A), while in *Apc^Δ580^* mice, the dominant products were prostaglandin E2 (PGE2) and 12-HETE (Figure 4A; Supplementary Figure 3A). Lipoxin A4 (LXA4) and LXB4 levels were also significantly higher in *Apc^Δ580^–15-LOX-1* than in *Apc^Δ580^*mice (Figure 4A). Levels of 15-LOX-1 enzymatic products (13-HODE, 15-HETE, LXA4, and LXB4) were significantly negatively correlated with tumor volumes in both *Apc^Δ580^* and *Apc^Δ580^–15-LOX-1* mice (Figures 4B-E). In contrast, levels of non–15-LOX-1 enzymatic products (PGE2, 5-HETE, 12-HETE, and LTB4) did not significantly correlate with tumor volumes (Supplementary Figures 3B-E).

**Figure 4.**
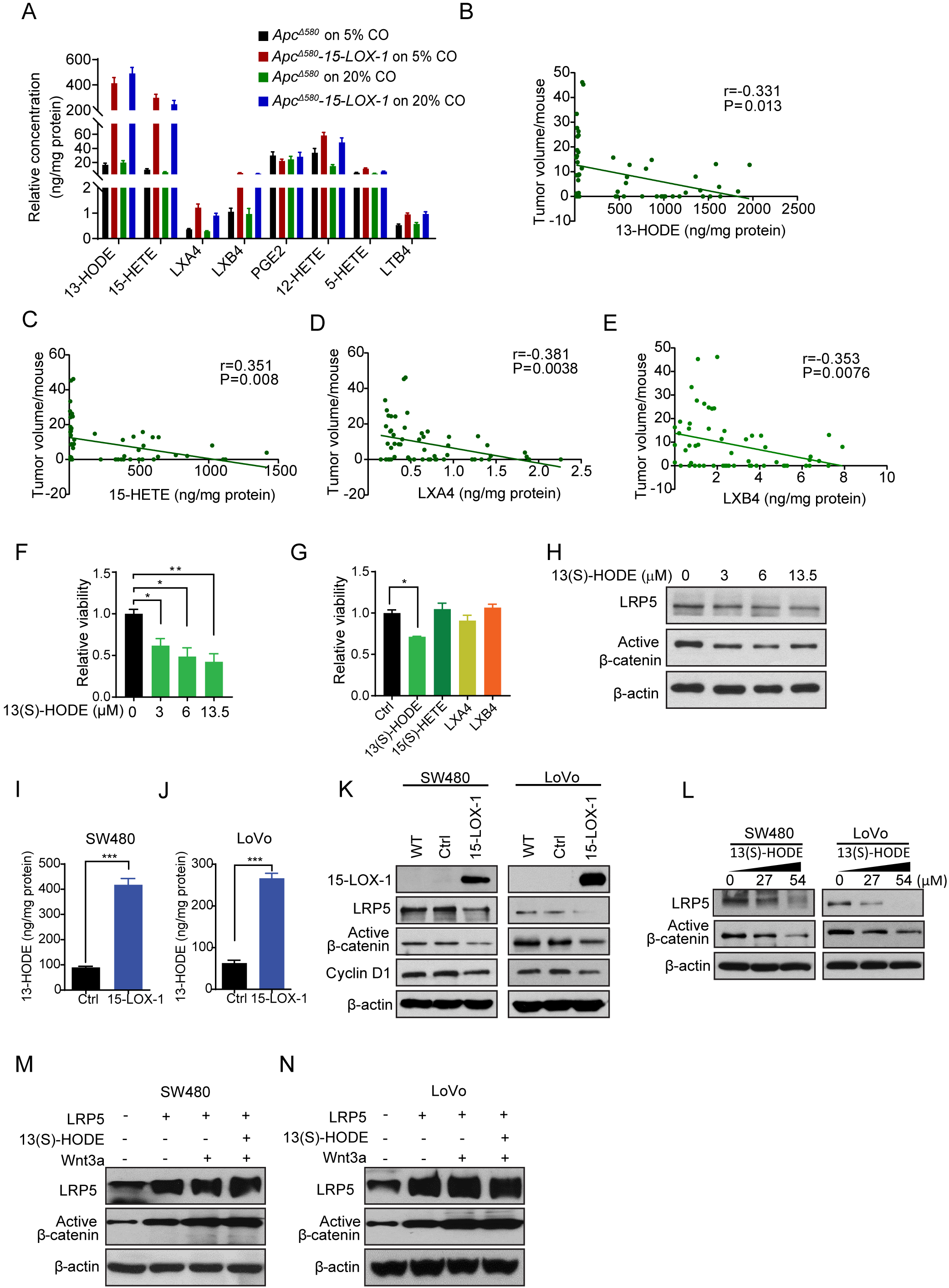
15-LOX-1 downregulates LRP5 expression via 13-HODE. **(A)** Eicosanoid metabolite profile of the colonic IECs from *Apc^Δ580^* and *Apc^Δ580^–15-LOX-1*, as described in Figures 1A-C, measured by liquid chromatography–tandem mass spectrometry (LC-MS/MS) (n=14 mice per group). **(B-E)** Spearman correlation analysis of colonic tumor volumes with levels of 13-HODE **(B)**, 15-HETE **(C)**, LXA4 **(D)**, or LXB4 **(E)** in IECs from the mice as described in **A** (n=56 for all 4 groups). **(F, G)** Cell viability of colonic organoids derived from IECs of *Apc^Δ580^*mice treated with 13(S)-HODE at 0, 3, 6, or 13.5 µM **(F)** or treated with 13(S)-HODE (3 µM), 15-HETE (2.15 µM), LXA4 (8.86 nM), or LXB4 (36.24 nM) for 6 days **(G)**, measured by CellTiter-Glo Luminescent Cell Viability Assay. **(H)** LRP5 and active β-catenin protein expression levels in organoid cells as described in panel **F**, treated with 13(S)-HODE at the indicated concentrations for 6 days. **(I, J)** Levels of 13-HODE in SW480 **(I)** and LoVo **(J)** cells stably transduced with either control (Ctrl) or 15-LOX-1 lentivirus, measured by LC-MS/MS. **(K)** 15-LOX-1, LRP5, and active β-catenin protein levels in SW480 and LoVo cells transduced with either Ctrl or 15-LOX-1 lentivirus, measured by Western blot. **(L)** LRP5 and active β-catenin protein levels measured by Western blot in SW480 and LoVo cells supplemented with 13(S)-HODE at the indicated concentrations in cell culture media for 48 hours. **(M, N)** LRP5 protein and active β-catenin protein levels in SW480 **(M)** and LoVo **(N)** cells transfected with control (-) or LRP5 expression vector (+) and treated with Wnt3a (100ng/ml) with or without 13(S)-HODE (27 µM) for 48 hours.

We then determined the mechanistic relevance of those correlations to our previously observed effects of 15-LOX-1 on stem cell regeneration (Figures 3D-F). 13(S)-HODE treatment inhibited regeneration of *Apc^Δ580^* IECs–derived organoids in a concentration-dependent manner (Figure 4F). In contrast, other 15-LOX-1 enzymatic products [15(S)-HETE, LXA4, and LXB4] had no effect on colonic organoid regeneration (Figure 4G) at concentrations proportional to their levels relative to 13-HODE levels in IECs of the *Apc^Δ580^* mice as measured by LC-MS/MS (Figure 4A). Notably, 13(S)-HODE treatment decreased LRP5 and active β-catenin protein levels in *Apc^Δ580^* IECs-derived organoids at the same 13(S)-HODE concentrations at which organoid regeneration was repressed (Figure 4H).

Next, we investigated the relevance of our mouse findings to human CRC. Re-expression of 15-LOX-1 in SW480 and LoVo human CRC cells via lentivirus infection significantly increased 13-HODE levels and inhibited LRP5 and active β-catenin protein levels (Figures 4I-K). Treatment of SW480 and LoVo with 13(S)-HODE downregulated LRP5 and active β-catenin protein levels (Figure 4L) at the same 13(S)-HODE concentrations that inhibited cell proliferation (Supplementary Figures 3F and G). In contrast, treatment with other 15-LOX-1 products such as 15(S)-HETE, LXA4 and LXB4 did not affect LRP5 and active β-catenin protein levels in both SW480 and LoVo cells (Supplementary Figures 3H and I). On examination of the mechanistic significance of LRP5 in these 13(S)-HODE effects, LRP5 overexpression reversed 13(S)-HODE’s downregulation of active β-catenin levels in SW480 (Figure 4M) and LoVo cells (Figure 4N). Together these results support the conclusion that 13-HODE mediates 15-LOX-1*–*induced downregulation of LRP5 expression.

### 15-LOX-1 regulates LRP5 expression in cell membrane

Membranous LRPs are tightly regulated at the protein stability level through balancing the fate of LRPs during their endocytosis with LDL between lysosomal degradation and recycling to the cell membrane (van Kerkhof et al., 2005). In testing 15-LOX-1’s effects on LRP5 protein stability, 15-LOX-1 re-expression in SW480 and LoVo cells decreased LRP5 protein stability following cycloheximide treatment (Figures 5A and B). Because LRP5 promotes 15-LOX-1 membranous translocation to oxidize LA-enriched LDL (Zhu et al., 2003), we investigated whether LA supplementation to CRC cells is sufficient to trigger 15-LOX-1 membranous re-localization and whether this re-localization affects LRP5 protein internalization and subsequently its membranous abundance. LA supplementation to LoVo and SW480 cells increased LRP5 re-distribution from the cell membrane to the cytoplasm within 2 hours when 15-LOX-1 was re-expressed in the cells, and this LRP5 re-distribution was associated with increased 15-LOX-1 protein distribution from the cytoplasm to the cell membrane (Figures 5C-F, Supplementary Figures 4A and B). To examine whether this observed re-distribution of LRP5 expression is due to an increased LRP5 internalization, we performed LRP5 internalization measurement using imaging flow cytometry with immunofluorescence-labeled LRP5 protein. LRP5 internalization was significantly higher in SW480 cells stably transduced with 15-LOX-1 lentivirus with LA exposure than in those transduced with control lentivirus with the same LA treatment (Figures 5G-I).

**Figure 5.**
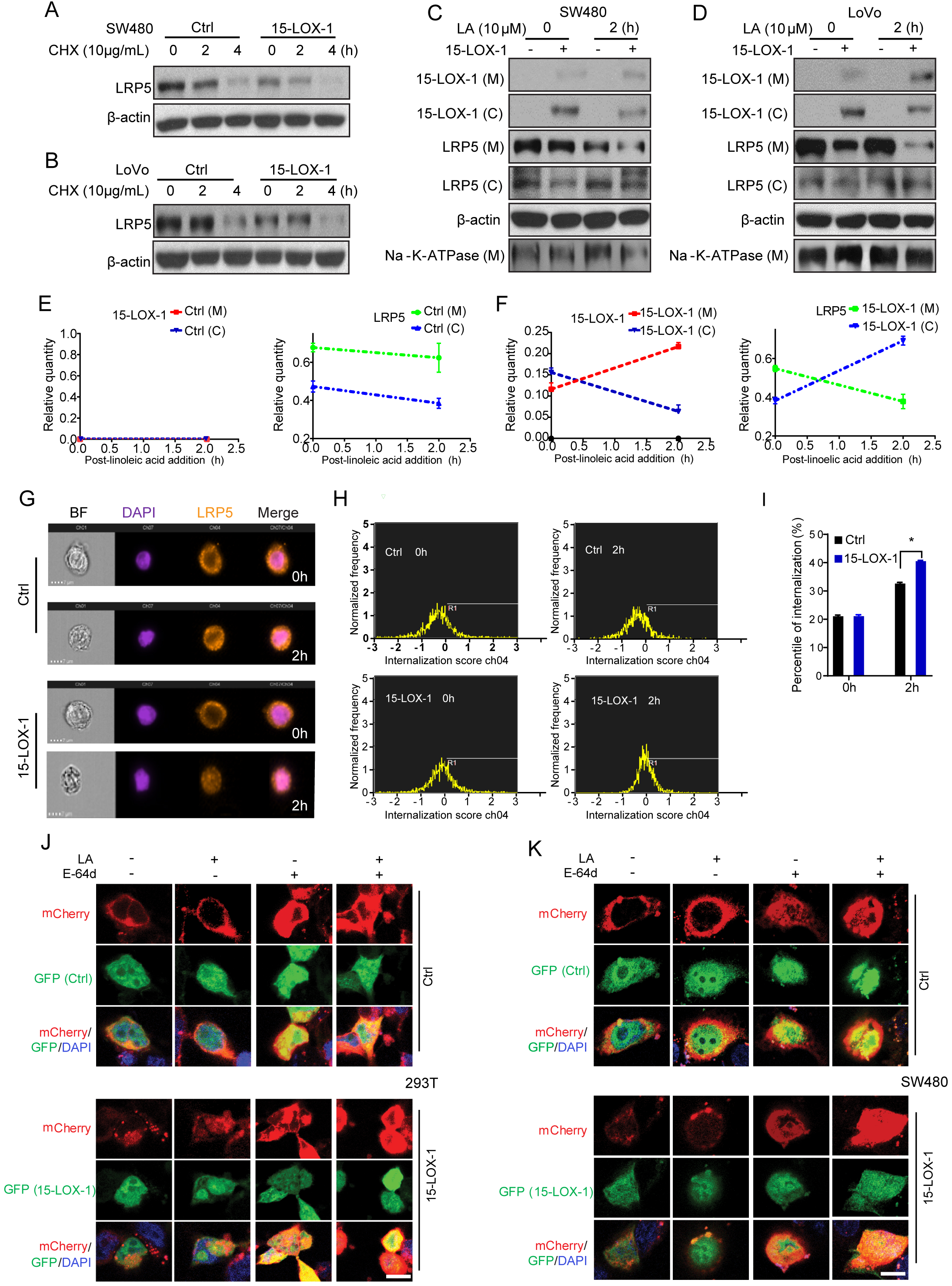
15-LOX-1 decreases cell membranous LRP5 levels. **(A, B)** LRP5 protein expression in SW480 **(A)** and LoVo **(B)** cells stably transduced with control (Ctrl) or 15-LOX-1 lentivirus and treated with 10 µg/mL cycloheximide (CHX) for 0, 2, and 4 hours. The whole-cell lysates were analyzed by Western blot. **(C, D)** Cytoplasmic (C) and membranous (M) LRP5 and 15-LOX-1 protein expression in SW480 **(C)** and LoVo **(D)** cells stably transduced with control (Ctrl) or 15-LOX-1 lentivirus and treated with 10 µM LA for 2 hours. The cells were processed into cytoplasmic and membranous protein fractions and then analyzed by Western blot. **(E, F)** The ratios of protein band densities of cytoplasmic 15-LOX-1 or LRP5 over β-actin and membranous 15-LOX-1 or LRP5 over Na-K-ATPase in Ctrl **(E)** or 15-LOX-1 **(F)** lentivirus transduced SW480 cells corresponding to panel **C**. **(G-I)** SW480 cells stably transduced with Ctrl or 15-LOX-1 lentivirus were treated with 10 µM LA for 2 hours. The cells were analyzed for LRP5 internalization by image flow cytometry assay. Representative images **(G, H)** and corresponding quantitative internalization percentile of LRP5 **(I)** for the indicated cells. **(J, K)** 293T **(J)** and SW480 **(K)** cells were co-transfected with mCherry-tagged LRP5 expression plasmid with pCMV5-GFP-IRES (GFP [Ctrl]) or with pCMV5-GFP-IRES-ALOX15 (GFP [15-LOX-1]) for 48 hours, then treated with E-64d (10 µM) or dissolvent (Ctrl) for 1 hour before cell exposure to LA (100 µM) supplement for another 4 hours. Representative microphotographs of LRP5 trans-localization traced with mCherry fluorescence protein by confocal microscope are shown. Scale bars: 2.5 µm (J) and 5.0 µm (K). BF: bright field.

To investigate whether this increased LRP5 internalization induced by 15-LOX-1 augments LRP5 lysosomal degradation, we transfected 293T cells with mCherry-labeled LRP5 expression vector to trace LRP5 distribution following LA supplementation, 15-LOX-1 re-expression, and treatment with E-64d, an irreversible broad-spectrum cysteine protease inhibitor that suppresses lysosomal protein degradation. LA supplementation increased membranous LRP5 protein levels while E-64d treatment increased both membranous and cytoplasmic LRP5 protein levels in the 293T cells transfected with GFP-labeled control vector. In contrast, transfection of GFP-labeled 15-LOX-1 vector in 293T cells reduced LRP5 levels, especially in the membranous compartment, while E-64d blocked those 15-LOX-1’s effects (Figure 5J). Similar results were obtained in SW480 cells (Figure 5K). These findings suggest that increased membranous internalization of LRP5 induced by 15-LOX-1 increases LRP5 lysosomal degradation.

### 15-LOX-1 inhibits SNX17-mediated LRP5 recycling to the cell membrane

Sorting nexin 17 (SNX17) binds membranous phosphatidylinositol 3-phosphate (PI3P) for anchorage to direct LRPs away from lysosomal degradation and recycle them into their membranous pools (van Kerkhof et al., 2005). We found that SNX17 overexpression increased LRP5 and active β-catenin protein levels in 293T and SW480 cells (Figure 6A). In turn, SNX17 downregulation by siRNA reduced LRP5 and active β-catenin protein levels (Figure 6B). Next, we used immunoprecipitation assays and immunoblotting to determine whether LRP5 binds to SNX17 to promote LRP5 protein recycling. When Myc-tagged LRP5 was expressed in 293T and SW480 cells, SNX17 was detected by immunoblotting in immunoprecipitated products with anti-c-Myc antibody (Figure 6C). Additionally, when Flag-tagged SNX17 was expressed in 293T and SW480 cells, LRP5 was also detected by immunoblotting in immunoprecipitated products with anti-Flag antibody (Figure 6D). These findings demonstrate the ability of LRP5 to bind SNX17. To examine whether SNX17 affects 15-LOX-1’s modulation of LRP5 protein stability, we performed a LRP5 cycloheximide pulse-chase assay with SNX17 and 15-LOX-1 overexpression. SNX17 overexpression reduced LRP5 degradation following cycloheximide treatment and, furthermore, blocked the augmentation of LRP5 degradation by 15-LOX-1 re-expression in SW480 cells without (Figures 6E) and with LA supplementation (Figure 6 F). We next examined whether 15-LOX-1 affects LRP5 or SNX17 binding to LA using a chemical ligation proximity assay. 15-LOX-1 re-expression markedly decreased the co-localization of alkynyl LA with SNX17 (Figure 6G) or with LRP5 (Figure 6H) in SW480 cells. In co-immunoprecipitation studies, 15-LOX-1 re-expression in SW480 cells with LA supplementation or treatment of SW480 cells with 13(S)-HODE inhibited LRP5 binding to SNX17 (Figure 6I), suggesting that 15-LOX-1 conversion of LA to 13(S)-HODE suppresses SNX17 binding to LRP5.

**Figure 6.**
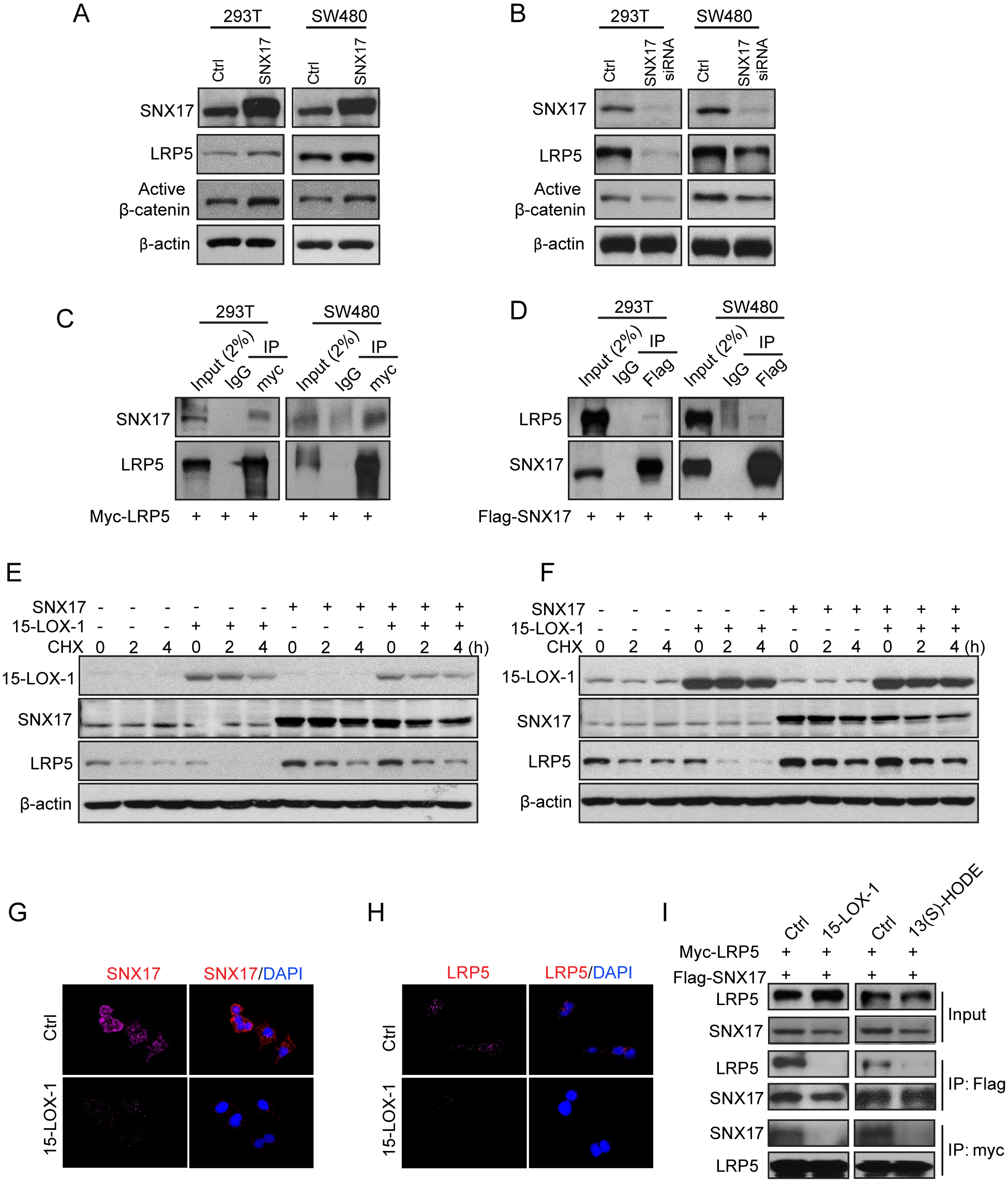
15-LOX-1 reduces membranous LRP5 levels by inhibiting SNX17-mediated LRP5 recycling to cell membrane. **(A)** SNX17, LRP5, and active β-catenin expression in 293T or SW480 cells transfected with control vector (Ctrl) or Flag-tagged SNX17 expression vector for 48 hours, then processed for Western blot. **(B)** SNX17, LRP5, and active β-catenin expression levels in 293T or SW480 cells transfected with control (Ctrl) or SNX17 siRNA pool for 72 hours, measured by Western blot. **(C)** SNX17 and LRP5 interaction assessment in 293T cells or SW480 cells transfected with Myc-tagged LRP5 expression vector for 48 hours. Cell lysates were immune-precipitated (IP) by anti-Myc antibody or rabbit immunoglobulin G (IgG) and then analyzed by immunoblotting with anti-SNX17 or anti-LRP5 antibody. **(D)** LRP5 and SNX17 interaction assessment in 293T cells or SW480 cells transfected with Flag-tagged SNX17 expression plasmid for 48 hours. Cell lysates were immune-precipitated by anti-Flag antibody or rabbit IgG and then analyzed by immunoblotting with anti-LRP5 or anti-SNX17 antibody. **(E, F)** SW480 cells stably transduced with control (-) or 15-LOX-1 (+) lentivirus were transfected with either SNX17 expression vector (+) or control vector (-) and treated with solvent **(E)** or 5 µM LA **(F)** for 48 hours, followed by treatment of cycloheximide (CHX) (10 µg/mL) for 0, 2, and 4 hours. SNX17, LRP5, and active β-catenin expression levels were measured by Western blot. **(G, H)** SW480 cells stably transduced with control (Ctrl) or 15-LOX-1 lentivirus were treated with 100 μM alkynyl LA for 36 hours, and the cells were then analyzed by proximity ligation assay using the Click chemistry reaction of alkynyl LA with SNX17 or with LRP5. Representative Click microphotographs for SNX17 **(G)** and LRP5 **(H)** are shown. **(I)** Effects of 15-LOX-1 or 13-HODE on LRP5 binding to SNX17. SW480 cells stably transduced with control (Ctrl) or 15-LOX-1 lentivirus were co-transfected with Myc-tagged LRP5 expression plasmid and Flag-tagged SNX17 expression plasmid for 48 hours, then treated with 10 µM LA for additional 24 hours (**left**). Wild-type SW480 cells were co-transfected with LRP5 and SNX17 expression plasmids for 48 hours, then treated with 27 µM 13(S)-HODE for another 24 hours (**right**). Cell lysates were immunoprecipitated by anti-Myc antibody or anti-Flag antibody and then analyzed by immunoblotting with anti-SNX17 or anti-LRP5 antibody. Scale bars: 50 µm (G and H).

### 15-LOX-1-mediated peroxidation of PI3P_LA to generate PI3P_13-HODE inhibits PI3P binding to SNX17 and thus suppresses LRP5 recycling to the cell membrane

Within endosomes, PI3P anchors SNX17 to recycle LRPs (van Kerkhof et al., 2005); we therefore examined the role of PI3Ps in LRP5 recycling by SNX17 using VPS34-IN-1. VPS34-IN-1 is a specific inhibitor of vacuolar protein sorting 34 (VPS34), which phosphorylates endosomal phosphatidylinositol to generate PI3Pthat recruits SNX17 recycling proteins such as LRP5 (van Kerkhof et al., 2005, Bago et al., 2014). VPS34-IN-1 decreased LRP5 protein expression, especially when 15-LOX-1 was re-expressed, in SW480 and LoVo cells (Figure 7A), suggesting that PI3P increases LRP5 protein expression and 15-LOX-1 might inhibit these PI3P effects. PI3P contains fatty acid residues that include LA, which could influence its biological functions (Gu et al., 2013). LA increased cellular PI3P production in SW480 cells, which was not altered by 15-LOX-1 re-expression (Figure 7B, Supplementary Figure 5A), suggesting that 15-LOX-1 downregulation of LRP5 was not related to alteration of overall PI3P abundance. On assessment of whether PI3P species were altered by 15-LOX-1, we measured PI3P_LA and PI3P_13-HODE species in IECs of *Apc^Δ580^* mice by liquid chromatography–high resolution mass spectrometry (LC-HRMS), and found that high dietary LA (20% vs. 5% corn oil) increased, while 15-LOX-1 transgenic expression in IECs decreased, PI3P(18:0/18:2)_LA levels (Figure 7C); in contrast, 15-LOX-1 transgenic expression in IECs considerably increased, PI3P (18:0/18:2OH)_13-HODE levels (Figure 7D). Similar changes were seen in the 18:1/18:2 species of PI3P (18:1/18:2)_LA (Figure 7E), PI3P (18:1/18:2 OH)_13-HODE (Figure 7F), and PI3P (16:0/18:2 OH)_13-HODE (Supplementary Figure 5B) in IECs of *Apc^Δ580^*and *Apc^Δ580^–15-LOX-1* mice.

**Figure 7.**
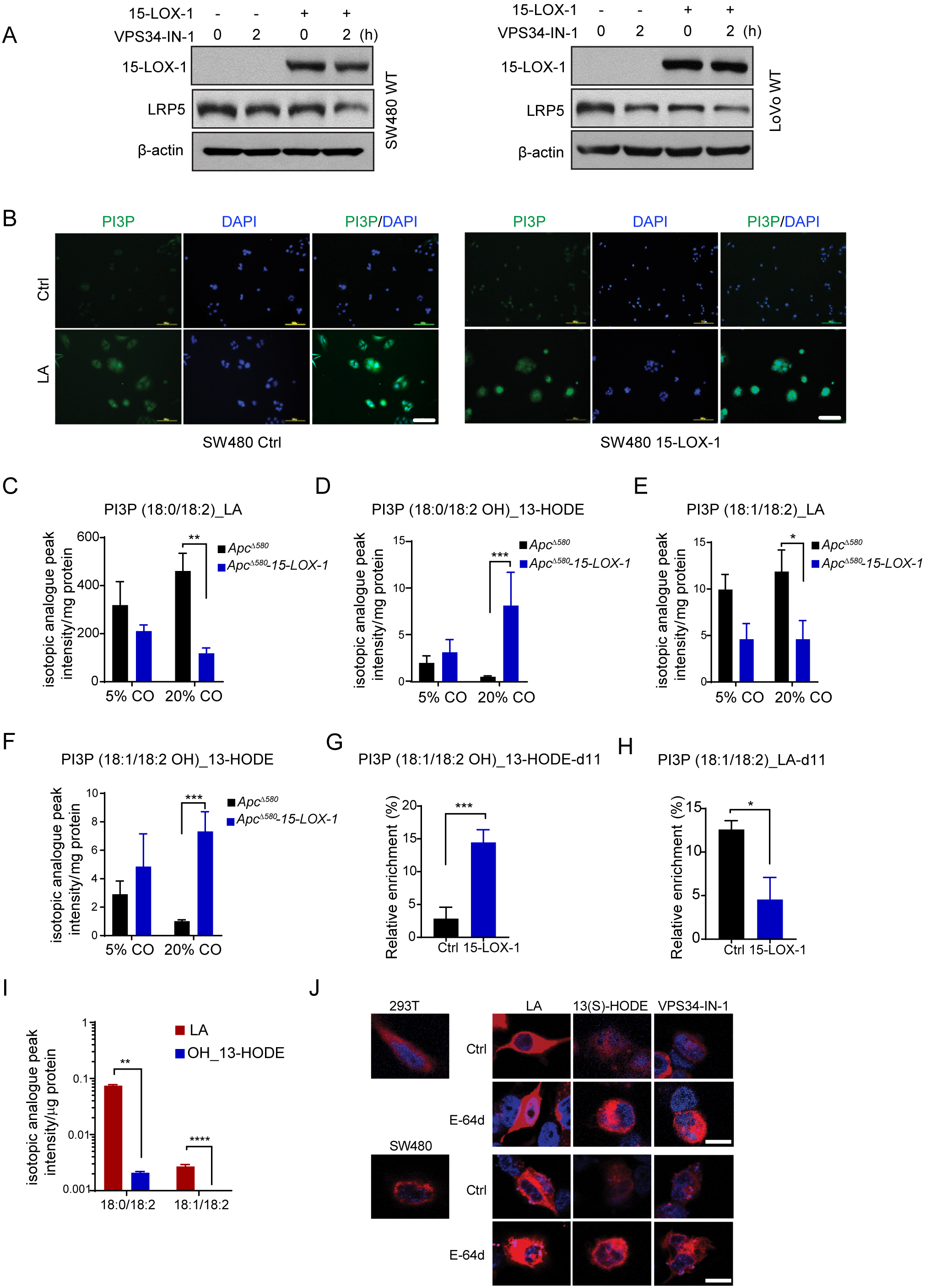
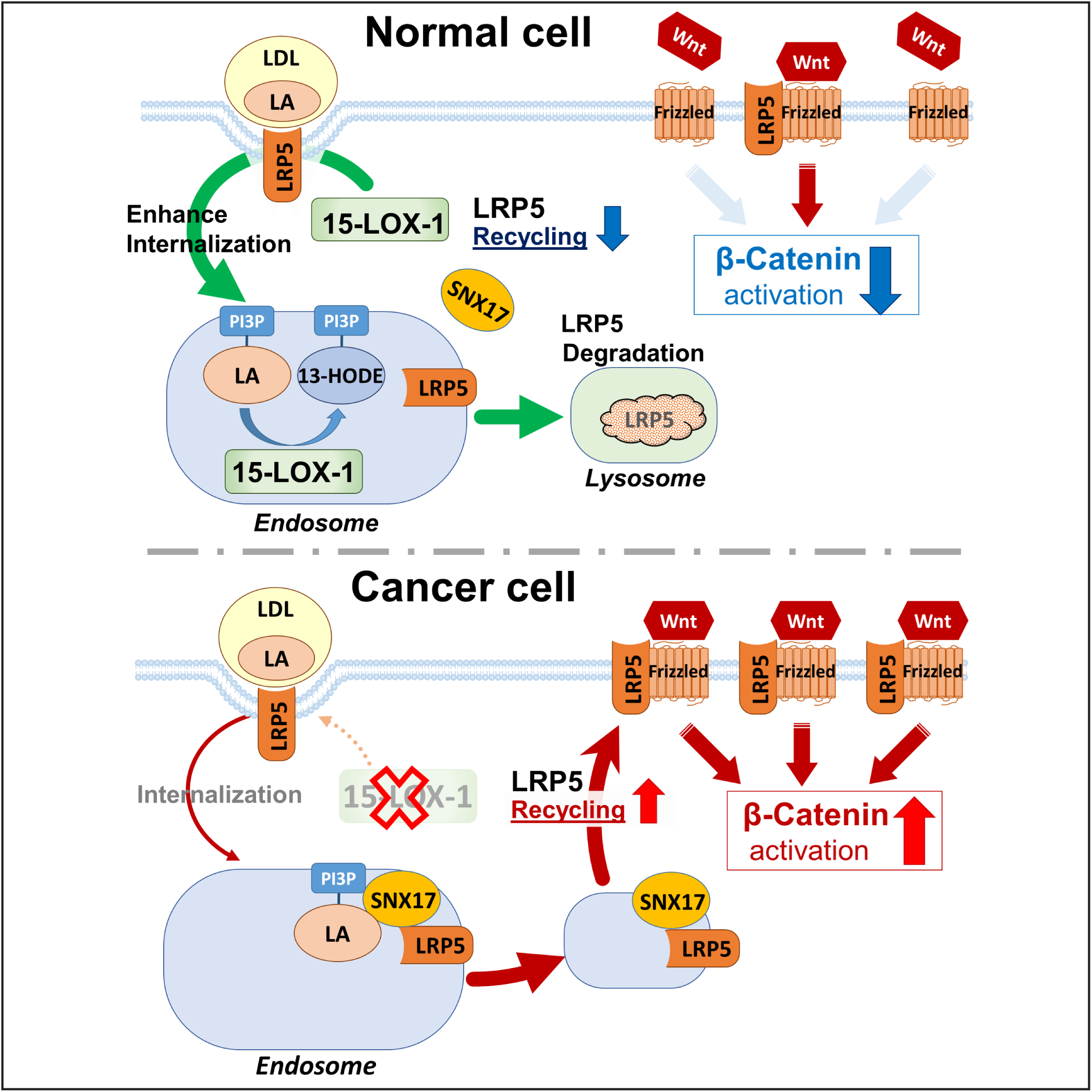
15-LOX-1 increases phosphatidylinositol 3-phosphate (PI3P)_13-HODE production, which inhibits PI3P’s binding to SNX17 in endosomes and subsequently suppresses LRP5 recycling to the cell membrane. **(A)** SW480 (**left**) or LoVo (**right**) cells stably transduced with control (Ctrl) or 15-LOX-1 lentivirus were treated with the PI3P biosynthesis inhibitor VPS34-IN-1 (1 µM) for 2 hours and processed for the indicated protein expression by Western blot. **(B)** SW480 cells stably transduced with control (Ctrl) or 15-LOX-1 lentivirus were treated with either vehicle solvent (Ctrl) or 5 µM LA for 48 hours. PI3P levels in those cells were analyzed by immunofluorescence staining. Representative microphotographs are shown. **(C-F)** *Apc^Δ580^* and *Apc^Δ580^–15-LOX-1* littermates at age 4 weeks fed either 5% or 20% corn oil were killed at age 14 weeks. The IECs of these mice were scraped and examined for PI3P incorporated with (18:0/18:2)_LA **(C)**, (18:0/18:2 OH)_13-HODE **(D)**, (18:1/18:2)_LA **(E)**, and (18:1/18:2 OH)_13-HODE **(F)** by LC-HRMS (n=3-4 mice per group). **(G, H)** SW480 cells stably transduced with control (Ctrl) or 15-LOX-1 lentivirus were treated with 100 µM LA-d11 for 36 hours in culture medium containing 5% dialyzed FBS. PI3P profiling was traced with d11 labeling by LC-HRMS. The percentage of PI3P (18:1/18:2)_13-HODE-d11 over total PI3P (18:1/18:2 OH)_13-HODE (**G**) and PI3P(18:1/18:2)_LA-d11 over total PI3P (18:1/18:2)_LA **(H)** were calculated as relative enrichment (%) and shown. **(I)** SW480 cells stably transduced with control (Ctrl) or 15-LOX-1 lentivirus were transfected with Flag-tagged SNX17 expression plasmid for 48 hours, followed by 100 µM LA treatment for another 24 hours. Cell lysates were immuno-precipitated by anti-Flag antibody–coated magnetic beads, then analyzed by LC-HRMS to examine the levels of PI3P incorporated with (18:1/18:2)_LA or with (18:1/18:2 OH)_13-HODE. **(J)** 293T (**top**) or SW480 cells (**bottom**) were transfected with mCherry-tagged LRP5 expression plasmid for 48 hours and treated with E-64d (10 µM) or dissolvent (Ctrl) for 1 hour, followed by LA (100 µM) or 13(S)-HODE (27 µM), or VPS34-IN-1 (1 µM) treatment for another 4 hours. Representative microphotographs of LRP5 trans-localization by confocal microscope are shown. Scale bars: 100 µm (B), 2.5 µm (J [top]), and 5 µm (J [bottom]).

To examine the human relevance of the 15-LOX-1 effect on PI3P species that contain LA, we performed LA-d11-metabolic tracing analyses in SW480 cells by LC-HRMS. We found that 15-LOX-1 re-expression considerably increased PI3P (18:1/18:2 OH)_13-HODE-d11 levels (Figure 7G) while decreasing PI3P (18:1/18:2)_LA-d11 levels (Figure 7H), but had less substantial effects on PI3P (18:0/18:2)_LA-d11 and PI3P (18:0/18:2 OH)_13-HODE-d11 production (Supplementary Figures 5C and D).

To elucidate whether PI3P_13-HODE has differential binding affinity to SNX17 compared to PI3P_LA, we examined by LC-HRMS the levels of PI3P species that were bound to immunoprecipitated Flag-tagged SNX17 protein ectopically over-expressed in SW480 cells with or without 15-LOX-1 re-expression. The relative enrichments of PI3P (18:0/18:2)_LA- and PI3P (18:1/18:2)_LA as measurements of their binding abilities to SNX17 were significantly higher than those of PI3P (18:0/18:2 OH)_13-HODE and PI3P (18:1/18:2 OH)_13-HODE (Figure 7I). These results show that conversion of (18:0/18:2)_LA or PI3P (18:1/18:2)_LA to PI3P (18:0/18:2 OH)_13-HODE or PI3P (18:1/18:2 OH)_13-HODE, significantly decreases PI3P binding to SNX17. Finally, given that PI3P-SNX17 plays a critical role in mediating LRP5 endosome trafficking and fate, we tracked LRP5 expression using mCherry red fluorescence in 293T and SW480 cells treated with LA or 13(S)-HODE. Compared with LA treatment, 13(S)-HODE treatment significantly decreased membranous LRP5 levels. This membranous LRP5 accumulation was enhanced by inhibiting lysosomal protein degradation via E-64d, but inhibited by VPS24-IN-1 (Figure 7J, Supplementary Figure 5E), demonstrating that 13(S)-HODE treatment promotes LRP5 trafficking for degradation rather than recycling to the cytoplasmic membrane. Together, these results demonstrate that 15-LOX-1 mediates conversion of PI3P_LA to PI3P_13-S-HODE, which decreases PI3P binding to SNX17 and, consequently, inhibits LRP5 recycling from lysosomes to the cell membrane.

## Discussion

We found that 1) 15-LOX-1 suppressed LA promotion of *Apc^Δ580^* mutation–induced CRC in mice; 2) 15-LOX-1 downregulated LRP5 expression and β-catenin activation in *Apc^Δ580^* mouse IECs, *Apc^Δ580^* IECs-derived organoid cells, human CRC cells, and human CRC–derived organoid cells; 3) 15-LOX-1 suppressed CRC stem cell self-renewal in mice and humans; 4) 13(S)-HODE, the primary 15-LOX-1 metabolic product, downregulated LRP5 expression and β-catenin activation in *Apc^Δ580^*IECs and human CRC cells; and 5) 15-LOX-1 promoted PI3P_LA conversion to PI3P_13-HODE, which markedly reduced PI3P binding to SNX17 and subsequently inhibited LRP5 recycling from lysosomal degradation to the cytoplasmic membrane pool. These findings identify a novel mechanism by which 15-LOX-1 regulates LRP5 protein recycling via its oxidative metabolism of LA residue within PI3P; this LRP5 modulation significantly affects Wnt/β-catenin signaling and its promotion of CRC.

High dietary LA promoted *Apc^Δ580^* mutation–driven CRC in mice, which was inhibited by IECs’ 15-LOX-1 transgenic expression. High dietary LA, which is typical of Western-type diets (Yang et al., 1996), has been reported to promote AOM-induced CRC in mice (Deschner et al., 1990). Our new findings show that LA promotion of CRC is not specific to CRC induction by AOM but also applies to other CRC experimental models, as in the case of an intestinally targeted *Apc^Δ580^* mutation, further experimentally confirming the potential deleterious effects of excess dietary LA. More importantly, these data identify a new mechanism by which LA promotes CRC by enhancing aberrant β-catenin activation and this activation can be strongly suppressed by its oxidative metabolism via 15-LOX-1. Our findings show the strong inhibiting effects of transgenic 15-LOX-1 expression in mouse IECs on *Apc* mutation–driven tumorigenesis using intestinally targeted *Apc^Δ580^* models that closely mimic human CRC (Hinoi et al., 2007). First, 15-LOX-1 transgenic expression in IECs significantly improved *Apc^Δ580^*mouse survival; after more than 1 year on a low LA diet, more than 50% of the mice with 15-LOX-1 transgenic expression remained alive, whereas none of the mice without 15-LOX-1 transgenic expression was alive. Second, 15-LOX-1 transgenic expression repressed especially large colorectal tumor formation, which is associated with CRC progression (Oshima et al., 2015), suggesting that 15-LOX-1 influences CRC beyond the initiation phase. Third, 15-LOX-1 repressed intestinal crypt proliferative zone expansion, an important mechanism by which β-catenin activation drives CRC tumorigenesis (van de Wetering et al., 2002). Finally, 15-LOX-1 repressed CRC stem cell self-renewal, a critical phenotypic feature by which CRC stem cells promote CRC progression. Thus, 15-LOX-1 suppresses LA-induced hyper-activation of Wnt/β-catenin signaling beyond *APC* mutations that can strongly promote CRC progression especially invasion^5^.

15-LOX-1 inhibited Wnt/β-catenin signaling via downregulation of LRP5. LRP5 is a key component of the LRP5/LRP6/Frizzled co-receptor group that acts as a Wnt receptor, which when activated stabilizes cytoplasmic β-catenin to potentiate its downstream signaling (MacDonald and He, Tortelote et al., 2017). LRP5 is also a member of the LRP family that functions as an endocytic receptor of LDL to mediate LDL internalization for intracellular processing via the endosomal/lysosomal system (Krieger and Herz, 1994, Herz et al., 2009, Tortelote et al., 2017). Cells tightly control LRPs membrane abundance, and subsequently Wnt signaling, via regulating the membranous abundance of LRPs by balancing their lysosomal degradation and recycling to the cell membrane through SNX proteins (e.g., SNX17) (van Kerkhof et al., 2005). LDL binding to LRPs has also been demonstrated to promote 15-LOX-1 translocation from the cytoplasm to the cell membrane to oxidize LA incorporated in complex lipids to form 13(S)-HODE (Zhu et al., 2003). Our results demonstrate for the first time that 15-LOX-1 translocation and oxidization of LA to 13-HODE impact LRP5 membranous abundance and subsequently Wnt/β-catenin signaling. LA upregulated LRP5 in CRC cells, which agrees with a previous report of LA upregulating LRP5 expression in peritoneal macrophages (Schumann et al., 2015). Our current findings however demonstrate for the first time that LA upregulates LRP5 to subsequently augment aberrant β-catenin activation beyond *APC* mutations in both *in-vivo* and *in-vitro* CRC models using both mouse models and human CRC cells. More importantly, 15-LOX-1 in IECs repressed aberrant catenin activation not only in the setting of inhibition of β-catenin signaling enhancement by excess dietary LA, but also even in the setting of low dietary LA intake.

15-LOX-1 downregulated LRP5 via 13-HODE. 15-LOX-1 transgenic expression in mouse IECs increased 13-HODE as the predominant oxidative metabolite compared to its other known metabolites of arachidonic acid (15-HETE, LXA4, and LXB4) (Brash, 1999, Serhan et al., 2008). While the levels of these various 15-LOX-1 products negatively correlated with the tumor burden in *Apc^Δ580^* mice, 13(S)-HODE was the only one among these metabolites to suppress *Apc^Δ580^*-mutant colonic stem cell self-renewal and to downregulate LRP5 and β-catenin activation. These findings indicate the specific mechanistic contribution of 13(S)-HODE to 15-LOX-1’s downregulation of LRP5 to suppress Wnt/β-catenin signaling, as a novel mechanism by which 15-LOX-1 suppresses CRC.

15-LOX-1 re-expression inhibited LRP5 protein recycling to cell membrane via PI3P_13-HODE. Our new data provide a novel mechanism for regulation of LRP5 membranous abundance and subsequently Wnt/β-catenin signaling via PI3P_13-HODE. These data show 15-LOX-1 recruitment to the membrane compartment increases LRP5 endosomal internalization and subsequently its lysosomal degradation. LRPs bind to SNX17; and SNX17 is anchored via PIPs, especially PI3P in early endosome, to promote subsequent LRPs recycling to the cell membrane (van Kerkhof et al., 2005). Our data demonstrate for the first time that modulation of LA as a PI3P lipid residue via 15-LOX-1 to form PI3P_13-HODE reduced SNX17 binding to LRP5, shifting LRP5 fate to lysosomal degradation and thus reducing its membranous recycling. 15-LOX-1 is unique among mammalian lipoxygenases in its ability to catalyze lipid peroxidation (e.g., LA) even when the substrate is bound by an ester linkage within a complex biological structure (Kuhn et al., 1990, Schewe et al., 1975). Our findings provide an example of the potential important role of this 15-LOX-1 function in important signaling events such as Wnt/β-catenin signaling. More importantly, this 15-LOX-1 regulatory mechanism is likely to play an important role in normal colonic cells to control LRP5 membranous abundance and to prevent excessive activation of Wnt/β-catenin signaling in normal cells that express 15-LOX-1. In contrast, cancer cells ubiquitously silence 15-LOX-1 expression (Moussalli et al., 2011, Il Lee et al., 2011), which could upregulate LRP5 especially in the presence of excess dietary LA to hyper-activate Wnt signaling/ β-catenin signaling in driving CRC via various pro-tumorigenesis mechanisms (e.g., stem cell self-renewal and colonic crypt proliferative zone expansion).

In summary, our findings demonstrate that increasing dietary LA intake promotes CRC by increasing Wnt/β-catenin activation via increasing LRP5 recycling to cell membrane, and that 15-LOX-1 represses LRP5 membranous recycling by converting PI3P_LA to PI3P_13-HODE to suppress Wnt/β-catenin signaling and subsequently CRC. This previously unknown regulation of Wnt/β-catenin signaling via 15-LOX-1 metabolism of PI3P_LA represents a modifiable mechanism, beyond *APC* mutations, by which Wnt/β-catenin signaling can be therapeutically targeted for CRC prevention and treatment.

## Author contributions

I.S. conceived the study. F.L., X.Z, Y.D., Y.L., Y.D., W.C., S.G., J.C.J. and L.A.V. performed various portions of animal experiments and histologic analyses and the *in-vitro* experiments. A.M. coordinated the collections of human colon cancer tissue samples. M.J.M. performed three-dimensional organoid culture. B.W. performed eicosanoid metabolite profiling by liquid chromatography–tandem mass spectrometry, and L.T. performed phosphatidylinositol 3-phosphate profiling by liquid chromatography–high resolution mass spectrometry. I.S., F.L., D.W., and X.Z. designed the experiments, analyzed data, and wrote the manuscript. D.W., P.Y., and P.L.L. provided conceptual feedback for the manuscript.

## Acknowledgments

We thank Dr.Jail Park at UT MD Anderson Cancer Center for his feedback on this work. We thank Ms. Sarah J. Bronson at the Department of Scientific Publications at MD Anderson Cancer Center for editing the article.

This study made use of the MD Anderson Cancer Center Genetically Engineered Mouse Facility, Functional Genomics Core, Flow Cytometry and Cellular Imaging Facility, Sequencing and Microarray Facility, and Research Animal Support Facility—Smithville Laboratory Animal Genetic Services, supported by Cancer Center Support Grant P30CA016672, and the MD Anderson Cancer Center metabolomics facility service, supported in part by Cancer Prevention Research Institute of Texas (CPRIT) grant RP130397 and NIH grant 1S10OD012304-01.

## STAR METHODS

### KEY RESOURCES TABLE

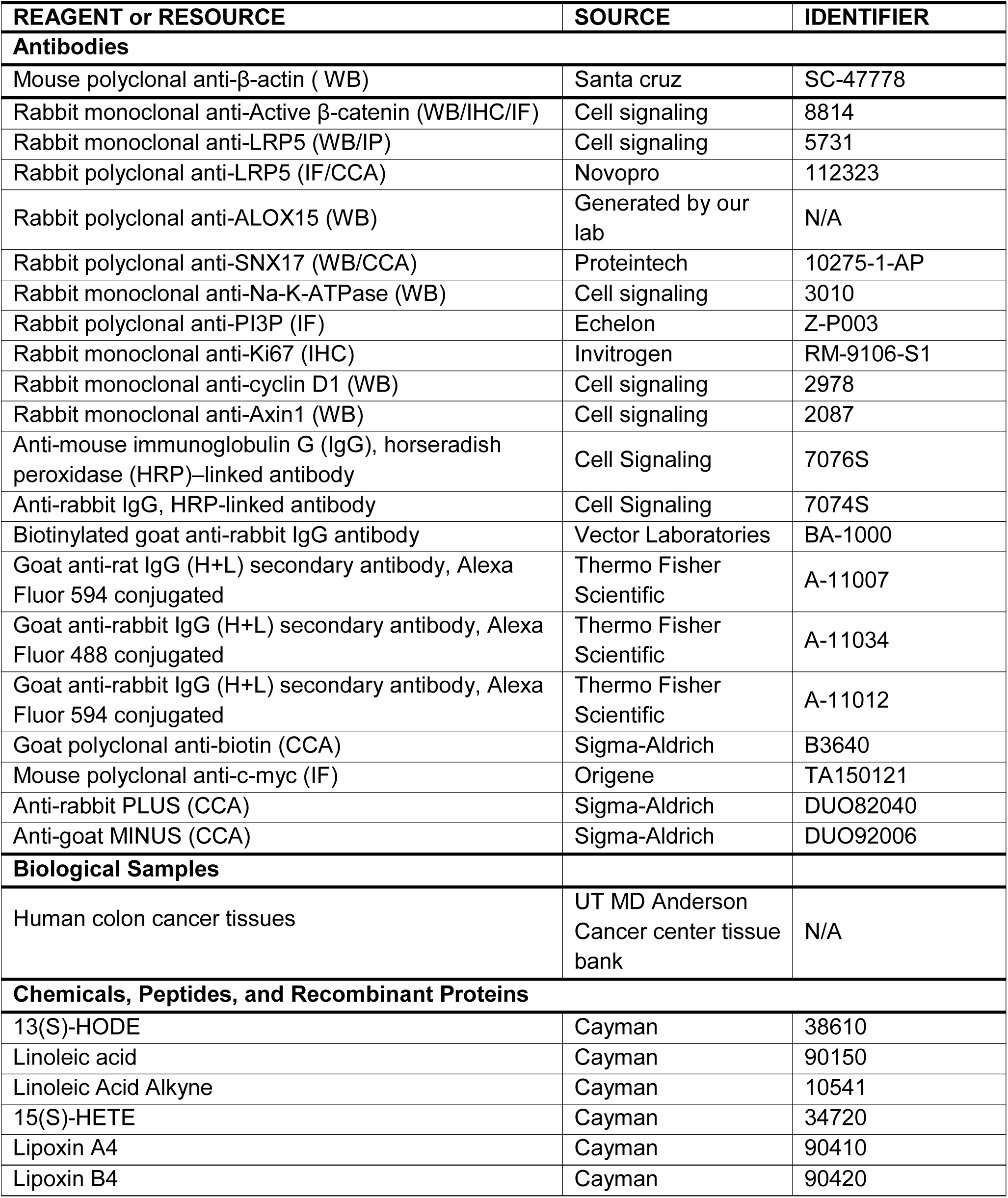
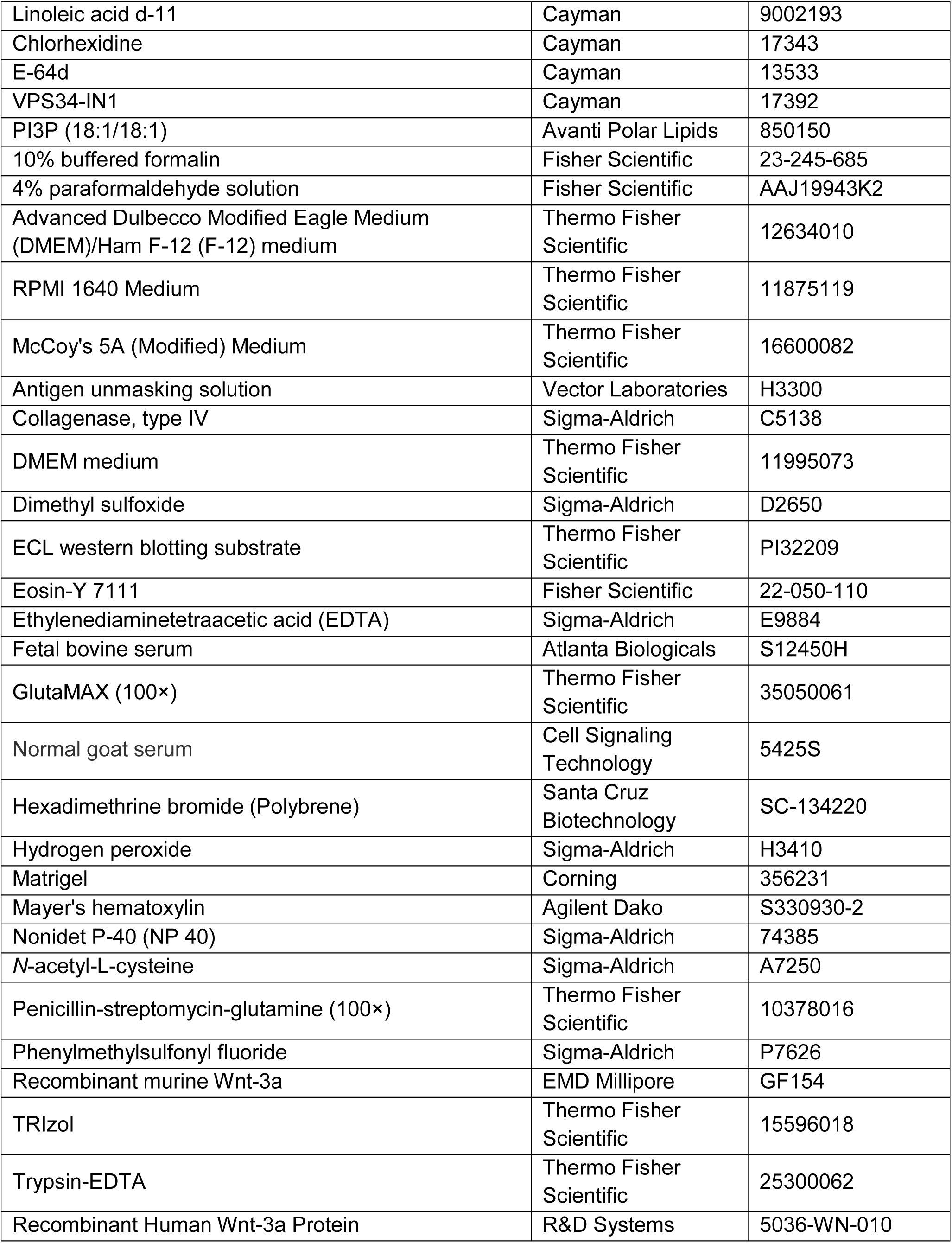
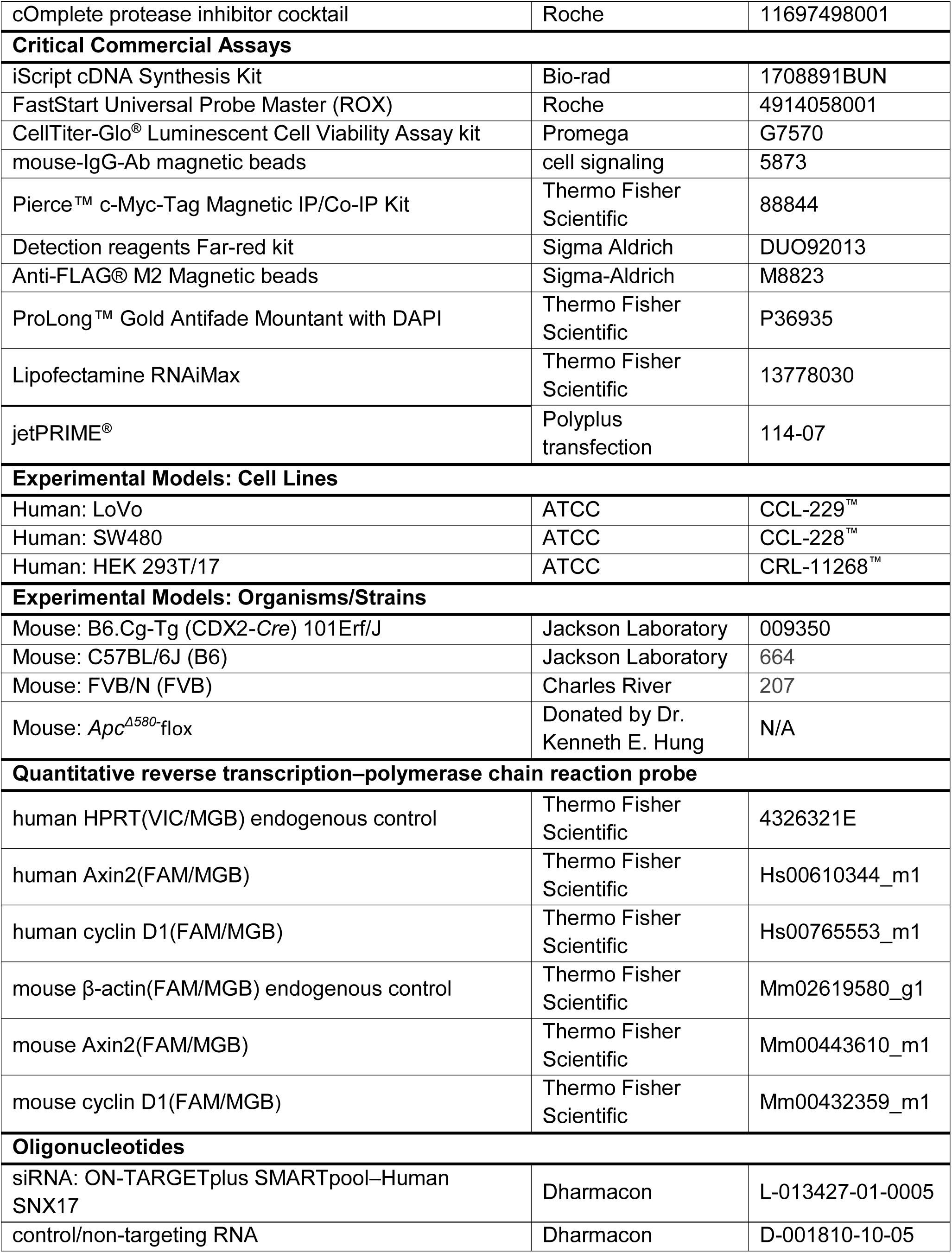
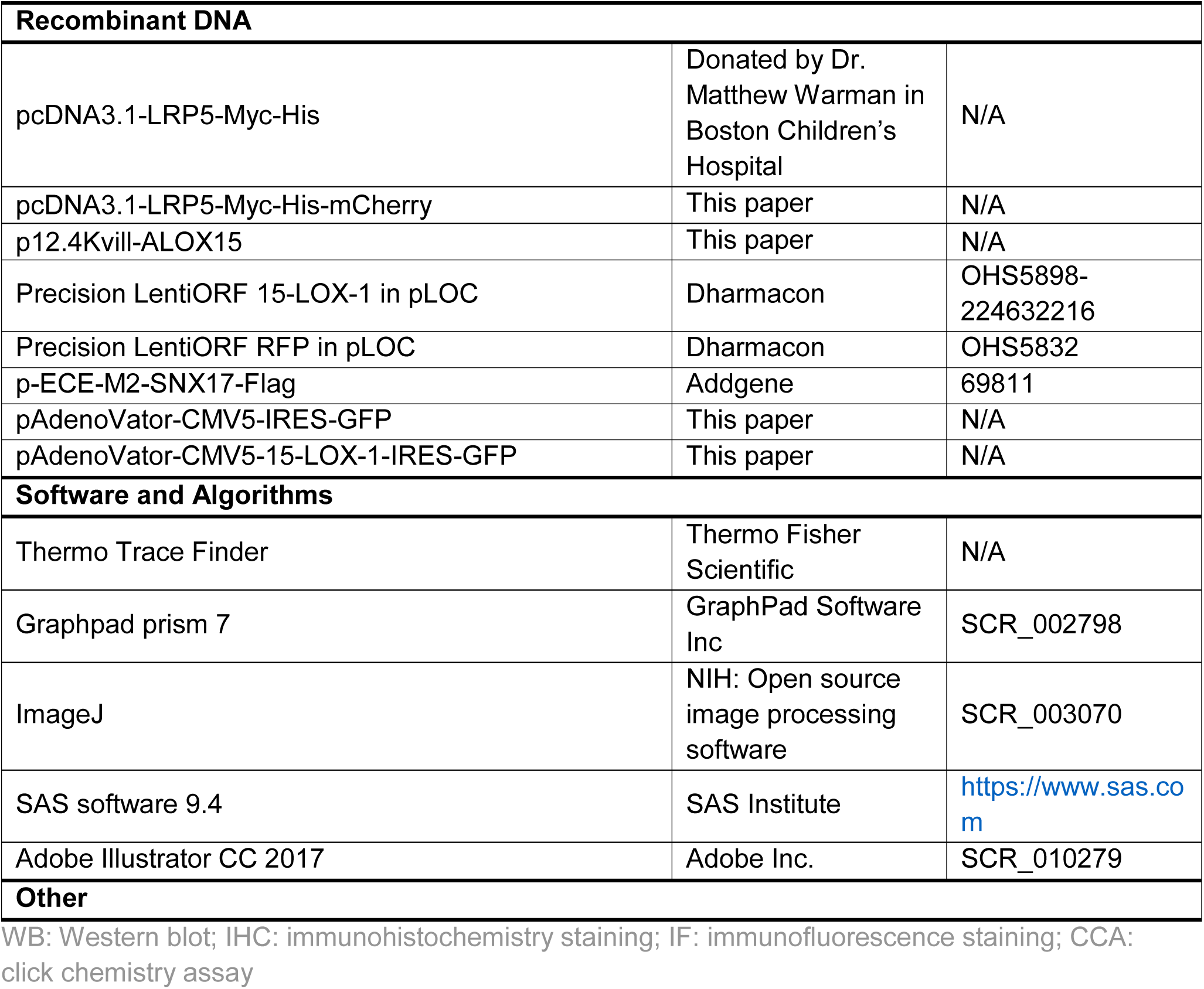

### CONTACT FOR REAGENT AND RESOURE SHARING

Further information and requests for resources and reagents should be directed to and will be fulfilled by the lead Contact, Imad Shureiqi.

### EXPERIMENTAL MODEL AND SUBJECT DETAILS

#### Cell culture

Human colon cancer cell lines SW480 and LoVo and human embryonic kidney cell line 293T were purchased from ATCC and were maintained in McCoy’s 5A, RPMI 1640 and Dulbecco modified Eagle (high glucose) media, respectively. All culture media were supplemented with 10% fetal bovine serum (FBS) and 1% penicillin/streptomycin, and all the cell lines were cultured in humidified atmosphere with 5% CO2 at 37°C. The cell lines were authenticated by short tandem repeat analyses and routinely tested for mycoplasma.

#### Animals

Mouse care and experimental protocols were approved by and conducted in accordance with the guidelines of the Animal Care and Use Committee of The University of Texas MD Anderson Cancer Center. We generated the mice using targeted 15-lipoxygenase-1 (*15-LOX-1*) overexpression in intestinal epithelial cells (IECs) via a villin promoter (designated as *15-LOX-1*-Gut mice) as described previously (Zuo et al., 2012). *B6.Cg-Tg (*CDX2*-Cre) 101Erf/J* (CDX2-*Cre*, #009350) mice were purchased from Jackson Laboratory*. Apc^Δ580^*-flox mice, in which *Apc* exon 14 is flanked with *loxP* sites, were a gift from Dr. Kenneth E. Hung at Tufts Medical Center (Hung et al., 2010). Breeding of *Apc^Δ580^*-flox mice with *Cre* recombinase–expressing mice (CDX2-*Cre*) deleted *Apc* exon 14 and consequently generated a codon 580 frame-shift mutation (*Apc^Δ580^*-flox; CDX2-*Cre*, designated as *Apc^Δ580^*mice) (Hung et al., 2010, Liu et al., 2019).

### METHOD DETAILS

#### Immunohistochemical analysis

Immunohistochemical staining was performed as described before (Zuo et al., 2017, Liu et al., 2019). Colon tissues from *Apc^Δ580^-15-LOX-1* mice or *Apc^Δ580^* WT littermates were fixed in 10% buffered formalin, embedded in paraffin, and cut into 5-µm sections. The tissue sections were deparaffinized and rehydrated, and antigen retrieval was performed with antigen unmasking solution (Vector Laboratories), and then the sections were incubated in blocking buffer (phosphate-buffered saline with 1.5% goat serum and 0.3% Triton X-100) for 1 hour at room temperature and incubated with primary antibodies in a humidified chamber at 4°C overnight. For immunohistochemistry staining (IHC), the following primary antibodies were used: active β-catenin (#8814; cell signaling); and Ki67 (#RM-9106-S1; Invitrogen, Carlsbad, CA). Subsequently, the tissue sections were incubated with biotinylated secondary antibodies (VECTASTAIN ABC kit; Vector Laboratories) for 1 hour, followed by incubation with avidin-coupled peroxidase (Vector Laboratories) for 30 minutes. 3, 3′-diaminobenzidine (DAB; Agilent Dako) was used as the chromogen, and the slides were counterstained with Mayer’s hematoxylin (Agilent Dako).

#### Cell immunofluorescence assay

The cells were seeded on coverslips in a six-well or twelve plate and cultured for 24 hours. Cells were fixed in cold 4% paraformaldehyde for 15 minutes and then simply rinsed with PBS. The cells were incubated with 0.1% Triton X-100 PBS on ice for 10 min with PBS, blocked in blocking buffer of 5% goat serum PBS containing 0.1% Triton X-100 at 37°C for 30 minutes, and then incubated with primary antibody at 4°C overnight. Subsequently, the cells were incubated with fluorochrome-conjugated secondary antibody at room temperature for 2 hours and then stained and mounted by ProLongGold Antifade Mountant with 4′, 6-diamidino-2-phenylindole (DAPI) (Thermo Fisher Scientific).

#### RNA isolation and quantitative real-time polymerase chain reaction (qRT-PCR) assay

Total RNAs from the cultured cells, IECs from colons, or three-dimensionally cultured colonic organoids were extracted using TRIzol reagent (Invitrogen), and then cDNA was synthesized using Bio-Rad cDNA Synthesis Kit (Bio-Rad Laboratories). qRT-PCR was performed using FastStart Universal Probe Master (ROX) (Roche) and the StepOnePlus PCR system (Applied Biosystems) as described before (Liu et al., 2019). An HPRT probe for human cells and β-actin probe for mouse cells were used as endogenous controls, and the mRNA expression levels of indicated targets were calculated using the comparative cycle threshold method (ddCt). All the probes were purchased from Applied Biosystems. All experiments were performed in triplicate.

#### Western blot

Western blot was performed as described previously (Zuo et al., 2017). Briefly, the cultured cells or the IECs scraped from the mice colons were lysed in the lysis buffer (Zuo et al., 2017) with sonication in ice water. The blots were probed with primary and secondary antibodies. The blots were then imaged using enhanced chemiluminescence (Thermo Fisher Scientific).

#### Eicosanoid metabolite profiling of IECs in the mice

Four-week-old *Apc^Δ580^* and *Apc^Δ580^*-15-LOX-1 littermates were fed diets containing either 5% or 20% corn oil (Envigo) for 10 consecutive weeks. The Mice (n=14 mice per group) were euthanized at age 14 weeks. The colon crypts were scraped for eicosanoid metabolite profiling by Liquid chromatography-tandem mass spectrometry measurements (LC-MS/MS) as described before (Shureiqi et al., 2010).

#### Generation of stable 15-LOX-1–overexpressing cell lines

Precision LentiORF 15-LOX-1 in pLOC plasmid (#OHS5898-224632216, clone ID: PLOHS_ccsbBEn_05805; Dharmacon), and Precision LentiORF RFP in pLOC vector (#OHS5832; Dharmacon) were packaged into lentivirus particles by MD Anderson’s shRNA and ORFeome Core Facility. LoVo and SW480 cells were transduced with lentiviral particles (10 MOIs) with 8 µg/mL hexadimethrine bromide. Twelve hours later, the culture medium was replaced with fresh medium containing blasticidin (15 µg/mL for SW480 cells, 20 µg/mL for LoVo cells). The medium was changed every 72 hours. The cells after blasticidin selection for 2 weeks were collected and expanded under blasticidin selection for further analyses.

#### Plasmid transient transfection

Flag-tagged SNX17 expression plasmid (pECE-M2-Snx17-Flag) was purchased from Addgene (#69811). Myc-tagged LRP5 expression plasmid (pcDNA3.1-Lrp5-myc-His) was a gift from Dr. Matthew Warman at Boston Children’s Hospital. pAdenoVator-CMV5-IRES-GFP and pAdenoVator-CMV5-15-LOX-1-IRES-GFP were obtained as described before (Shureiqi et al., 2003). To insert the mCherry coding sequence into the C-terminus of this Lrp5 expression plasmid, a fusion PCR method was performed as follows: a pair of primers (Lrp5-F: 5′-CTCCCTCGAGACCAATAACAAC-3′; Lrp5-R: 5′-CTCGCCCTTGCTCACGGATGAGTCCGTGCAGGGGG-3′) were used to amplify the C-terminus of Lrp5, and another pair of primers (mCherry-F: 5′-CTGCACGGACTCATCCGTGAGCAAGGGCGAGGAGGA-3′; mCherry-R: 5′-CCAGTTTAAACTTACTTGTACAGCTCGTCCATGCC-3′) were used to amplify mCherry cDNA. The fusion PCR products were then amplified by primers Lrp5-F and mCherry-R using C-terminus of Lrp5 and mCherry cDNA fragments as the PCR templates. The fusion PCR products were constructed into pcDNA3.1-Lrp5-Myc-His using PmeI and XhoI restriction sites.

#### siRNA transient transfection

Cells were seeded and cultured to 60% confluence and then transiently transfected with 100 nM ON-TARGETplus siRNA SMARTpool (human SNX17 [Dharmacon #L-013427-01-0005] or control/non-targeting [Dharmacon #D-001810-10-05]) using Lipofectamine RNAiMax (#13778030, Thermo Fisher Scientific) transfection reagent following the manufacturer’s protocol. The cells were further analyzed 48-72hours after transfection.

#### Cell proliferation and viability assay

The cells were seeded into a 96-well plate (3×10^3^/well). Twenty-four hours later, the cells were treated with 13(S)-HODE (#38610, Cayman) or dissolvent in 5% dialyzed FBS of the medium. The cell proliferation and viability were analyzed at 0, 24, 48, and 72 hours after treatment unless otherwise specified using MTT assay.

#### 15-LOX-1 and LRP5 translocation analysis

SW480 and LoVo cells stably transduced with 15-LOX-1 or control lentivirus were seeded in a six-well plate (5×10^5^/well). Twenty-four hours later, the cells were cultured in serum-free medium for another 24 hours, followed by treatment with 10 µM LA dissolved in 5% dialyzed FBS of the medium for 2 hours. Cells were lysated in chilled lysis buffer (50 mM HEPES [4-(2-hydroxyethyl)-1-piperazineethanesulfonic acid], 150 mM NaCl, 3 mM MgCl_2_ in PBS, pH 7.5) with proteinase inhibitor cocktail (#11697498001; Roche). After the pellets were discarded by spinning down at 1000*g* for 3 minutes, the supernatants were centrifuged at 4°C at 21,460*g* for 1 hour. The supernatants/cytoplasmic proteins were added with a final 1% sodium dodecyl sulfate, and membrane/cytoskeleton protein pellets were then re-suspended with 1% sodium dodecyl sulfate in the same lysis buffer as mentioned above. Cytoplasmic and cell membranous 15-LOX-1 and LRP5 expression levels were measured by Western blot.

#### Organoid culture for mouse colon epithelial cells and human CRC tissues and organoid cell viability assay

Six-week-old *Apc^Δ580^* and *Apc^Δ580^–15LOX-1* littermates (n=6 per group) were killed, and their colons were harvested and digested for three-dimensional organoid culture using the method described by Clevers and colleagues with some modifications (Sato et al., 2011, Liu et al., 2019). Human CRC tissues for three-dimensional organoid culture were obtained from CRC surgery from patients at MD Anderson Cancer Center with Institutional Review Board approval. Briefly, the glands were isolated with 10 mM EDTA by incubation for 10 minutes at room temperature. Equal amounts of glands (500 glands/well) were seeded in growth factor–reduced Matrigel (Corning) with the organoid culture medium (Sato et al., 2011) in 24 wells of plates. Mouse primary organoid cells (P0) were digested into single cells with TrypLE (Thermo Fisher Scientific), seeded onto Matrigel, and cultured using the same method as described above. On day 8 of the culture, primary (P0) and secondary (P1) organoids were imaged, counted, and harvested for further analyses. In addition, mouse organoid cells were treated with 13(S)-HODE (0 µM, 3 µM, 6 µM, and 13.5 µM), 15-HETE (2.15 µM), LXA4 (8.86 nM), and LXB4 (36.24 nM) in the organoid culture medium, and the medium was changed every 3 days. On day 7 of treatment, organoid cell viability was measured using CellTiter-Glo Luminescent Cell Viability Assay kit (Promega) according to the manufacturer’s manual.

Human CRC organoids were passaged at least five generations and then digested into single cells with TrypLE (Thermo Fisher Scientific) and plated in 24-well plates with the Ultra-Low attachment surface (Corning). The cells were then transduced with 15-LOX-1 or control lentivirus particles. Twelve hours after lentivirus transduction, the organoid cells were seeded onto Matrigel and cultured using the same method as described above. On day 7 of culture, the organoids transduced with 15-LOX-1 or control lentivirus were imaged, counted, and harvested for further analyses.

#### Image flow cytometry for LRP5 internalization analysis

SW480 cells stably transduced with 15-LOX-1 or control lentivirus were seeded in 6-cm dishes (1×10^6^ per dish). Twenty-four hours later, the cells were starved in a culture of serum-free medium for another 24 hours, then treated with 10 µM LA in 5% dialyzed FBS of the medium for 2 hours. Then, the cells were digested and fixed in 4% paraformaldehyde on ice for 15 minutes, followed by incubation with blocking buffer with 0.1% Triton X-100 at room temperature for 1 hour. Subsequently, cells were incubated with anti-LRP5 primary antibody (1:200, Novopro) for 1 hour and Alexa Fluor 594 secondary antibody (1:250, #A-11012; Invitrogen) for 30 minutes at room temperature. Before the analysis, the cells were stained with DAPI (5 μg/mL) (#D9542, Sigma-Aldrich) for 5 minutes. LRP5 internalization analysis was performed using Amnis FlowSight and ImageStreamX (LUMINEX) according to the manufacturer’s instructions.

#### Immunoprecipitation and co-immunoprecipitation assays

293T cells or SW480 cells were seeded in six-well plates (5×10^5^/well). Twenty-four hours later, the cells were transiently transfected with Flag-tagged SNX17 expression plasmid (pECE-M2-Snx17-Flag), Myc-tagged LRP5 expression plasmid (pcDNA3.1-Lrp5-myc-His), or their control plasmids using jetPRIME transfection reagent (#114-07, Polyplus-transfection) and were harvested 48 hours after transfection for immunoprecipitation or co-immunoprecipitation assays. These assays were performed using anti-Flag M2-tagged magnetic beads (#M8823; Sigma-Aldrich) and/or Pierce c-Myc–tagged magnetic IP/Co-IP Kit (#88844; Thermo Fisher Scientific) according to the manufacturer’s protocol instructions. The same amount of the lysates was incubated with magnetic bead–conjugated mouse immunoglobulin G (#5873, Cell Signaling Technology) as a control. The subsequently precipitated proteins were immunoblotted using the indicated antibodies as described in the corresponding Figure legends.

#### Click chemistry assay

A click chemistry assay was performed as described previously (Gao and Hannoush, 2016) except LA alkyne (Alk-LA) was used instead of Alk-C16 and anti-SNX17 or anti-LRP5 antibody was used instead of anti-Wnt-3a antibody. Briefly, SW480 cells stably transduced with 15-LOX-1 or control lentivirus were seeded on micro cover slips (#10026-136, VWR) in a 12-well plate and cultured in 10% FBS of McCoy’s 5A medium. The cells were treated with 100 μM Alk-LA for 36 hours. Then, the cells were fixed with pre-chilled methanol at −20°C for 10 minutes, followed by incubation with 0.1 % Triton X-100 PBS for 5 minutes at room temperature. The mixture solution of 5 mM biotin-azide, 50 mM Tris (2-carboxyethyl) phosphine hydrochloride (Sigma-Aldrich), and 50 mM CuSO_4_ in PBS was added to the cells and incubated at room temperature for 1 hour. The cells were then incubated with blocking buffer (5 % BSA and 0.3 % Triton X-100 in PBS) at room temperature for 1 hour, probed with either anti-SNX17 antibody (1:200; Proteintech) or anti-LRP5 antibody (1:200; NovoPro) at 4°C overnight, then probed with anti-biotin antibody (1:3000, #ab36406; Abcam) at room temperature for 1 hour, and then incubated with the oligonucleotide-conjugated secondary antibody solution by mixing Duolink In Situ PLA Probe Anti-Rabbit PLUS and Anti-Goat MINUS (#DUO92002 and #DUO92006, respectively; Sigma-Aldrich) together in blocking buffer at 37°C for 1 hour. Finally, the cells were incubated with ligation solution (#DUO92013; Sigma-Aldrich) in water, then with amplification solution (#DUO92013, Sigma-Aldrich) at 37°C for 100 minutes in a dark area and then mounted with ProLong Gold Antifade Mountant with DAPI (Thermo Fisher Scientific).

#### Cycloheximide treatment for protein degradation analysis

SW480 and LoVo cells stably transduced with 15-LOX-1 and control lentivirus were seeded in six-well plates (6×10^5^ per well). The cells were starved with serum-free medium for another 24 hours before 10 µg/mL cycloheximide (#14126, Cayman Chemical) dissolved in DMSO was added. The cells were harvested at 0, 2, and 4 hours after treatment for further analyses by Western blot.

#### In vitro PI3-kinase VPS34 inhibition assay

SW480 and LoVo cells stably transduced with 15-LOX-1 and control lentivirus were seeded in 6-well plates (3×10^5^ per well). 24 hours later, the cells were treated with 10 µM LA in 5% FBS of the cell culture medium or an equal amount of dissolvent (DMSO) for 48 hours, then treated with 1 µM class III phosphoinositide 3-kinase vacuolar protein sorting 34 Inhibitor 1 (VPS34-IN-1) in serum-free medium for 2 hours. Then, the cells were harvested and measured for LRP5 expression by Western blot.

#### Analysis of phosphatidylinositol 3-phosphate (PI3P) profile by liquid chromatography– high resolution mass spectrometry (LC–HRMS)

The following experiments were performed for LC–HRMS: 1) Colonic epithelial cells were scraped from *Apc^Δ580^* and *Apc^Δ580^-15-LOX-1* mice at 4 weeks fed with diets containing either 5% or 20% corn oil for 10 weeks (n=3-4 mice per group). 2) For the tracing-combined assay, SW480 cells stably transduced with 15-LOX-1 and control lentivirus were seeded in 10-cm dishes to reach 80% confluence, then treated with 100 μM LA–d11 (Cayman Chemical) in 5% dialyzed FBS of McCoy’s 5A medium for 24 hours. The cells were quickly washed with pre-chilled PBS to remove medium completely and harvested. 3) For the immunoprecipitation-combined assay, SW480 cells stably transduced with 15-LOX-1 and control lentivirus were seeded in 10-cm dishes and transfected with Flag-tagged SNX17 expression plasmid (pECE-M2-Snx17-Flag) using jetPRIME for 48 hours. The cells were then treated in 100 μM LA for another 24 hours and harvested for processing immunoprecipitation using anti-Flag M2 Magnetic beads. Then, precipitated samples were collected.

The samples collected from three experiments described above were extracted for lipid metabolites using cold chloroform/methanol/water acidified with hydrochloric acid (Kim et al., 2010). The extracted samples were centrifuged at 4°C at 17,000*g* for 5 minutes, and organic layers were transferred to clean tubes followed by evaporation to dryness under nitrogen. Samples were reconstituted in 90/10 acetonitrile/water and injected into a Thermo Fisher Scientific Vanquish liquid chromatography system containing a SeQuant ZIC-pHILIC 4.6×100 mm column with a 5-µm particle size. Mobile phase A (MPA) was acetonitrile containing 95/5 of acetonitrile/200mM ammonium acetate, pH5.8. Mobile phase B (MPB) was 90/5/5 of water/acetonitrile/200mM ammonium acetate, pH5.8. The flow rate was 300 µL/minutes at 35°C, and the gradient conditions were initially 15% MPB, increased to 95% MPB at 10 minutes, held at 95% MPB for 5 minutes, and returned to initial conditions and equilibrated for 5 minutes. The total run time was 20 minutes. Data were acquired using a Thermo Fisher Scientific Orbitrap Fusion Tribrid mass spectrometer with electrospray ionization negative mode at a resolution of 240,000. Then, the raw files were imported to TraceFinder software for final analysis (Thermo Fisher Scientific). The fractional abundance of each isotopologue is calculated by the peak area of the corresponding isotopologue normalized by the sum of all isotopologue areas. The isotopic relative enrichments of PI3Ps incorporated with LA or OH_13-HODE were calculated by comparing the peak areas normalized by the standard concentration of PI3P (18:1/18:1) and protein levels of each sample.

#### Mouse intestinal tumorigenesis evaluation and survival experiments

*Apc^Δ580^* mice were bred with *15-LOX-1*-Gut mice to produce *Apc^Δ580^*; *15-LOX-1*-Gut mice, designated as *Apc^Δ580^*–*15-LOX-1* mice. Four-week-old *Apc^Δ580^*and *Apc^Δ580^*-*15-LOX-1* littermates were fed diets containing either 5% or 20% corn oil (Envigo) for 10 consecutive weeks (n=14 mice per group). Mice were euthanized at age 14 weeks. The colons from the rectum to the cecum were removed and washed with phosphate-buffered saline (PBS) and photographed, tumors were counted, and tumor sizes were measured. The tumor volumes were calculated using the formula V (volume)=(length ×width^2^)/2 (Szot et al., 2018). The distal one third of colons were collected in 10% neutral formalin for hematoxylin and eosin staining and immunohistochemical staining (IHC) and the other two thirds of colons were scraped to harvest colonic epithelial cells for further analyses (e.g. RNA, protein, and mass spectrometry) as described in the corresponding supplementary method sections.

Survival experiments were also performed for *Apc^Δ580^*and *Apc^Δ580^*–*15-LOX-1* mice (n=14-20 mice per group) on a diet containing either 5% or 20% corn oil. The mice were followed until they required euthanasia on the basis of at least one of the following preset criteria: (1) persistent rectal bleeding for 3 consecutive days or/and (2) weight loss of more than 20%.

#### Statistical analysis

Statistical significance was determined by the unpaired Student *t*-test, chi-square test, or analysis of variance (one-way or two-way) with Bonferroni adjustments for all multiple comparisons. The significance of correlations was determined by Spearman correlation coefficient test. The survival time was calculated by the Kaplan-Meier method and compared between groups using the log-rank test. All tests were two-sided, and significance was defined as *P*<0.05. Data were analyzed using SAS software, version 9.4 (SAS Institute) or GraphPad Prism 7.01 (GraphPad Software). The data are presented as mean ± standard deviation or mean ± standard error (\**P*<0.05, \*\**P*<0.01, \*\*\**P*<0.001, and \*\*\*\**P*<0.0001).

