## Supplemental figure1-5 and legend for "Suppression of membranous LRP5 recycling, WNT/β-catenin signaling, and colorectal tumorigenesis by 15-LOX-1 peroxidation of PI3P_linoleic acid"

Supplementary Figure 1

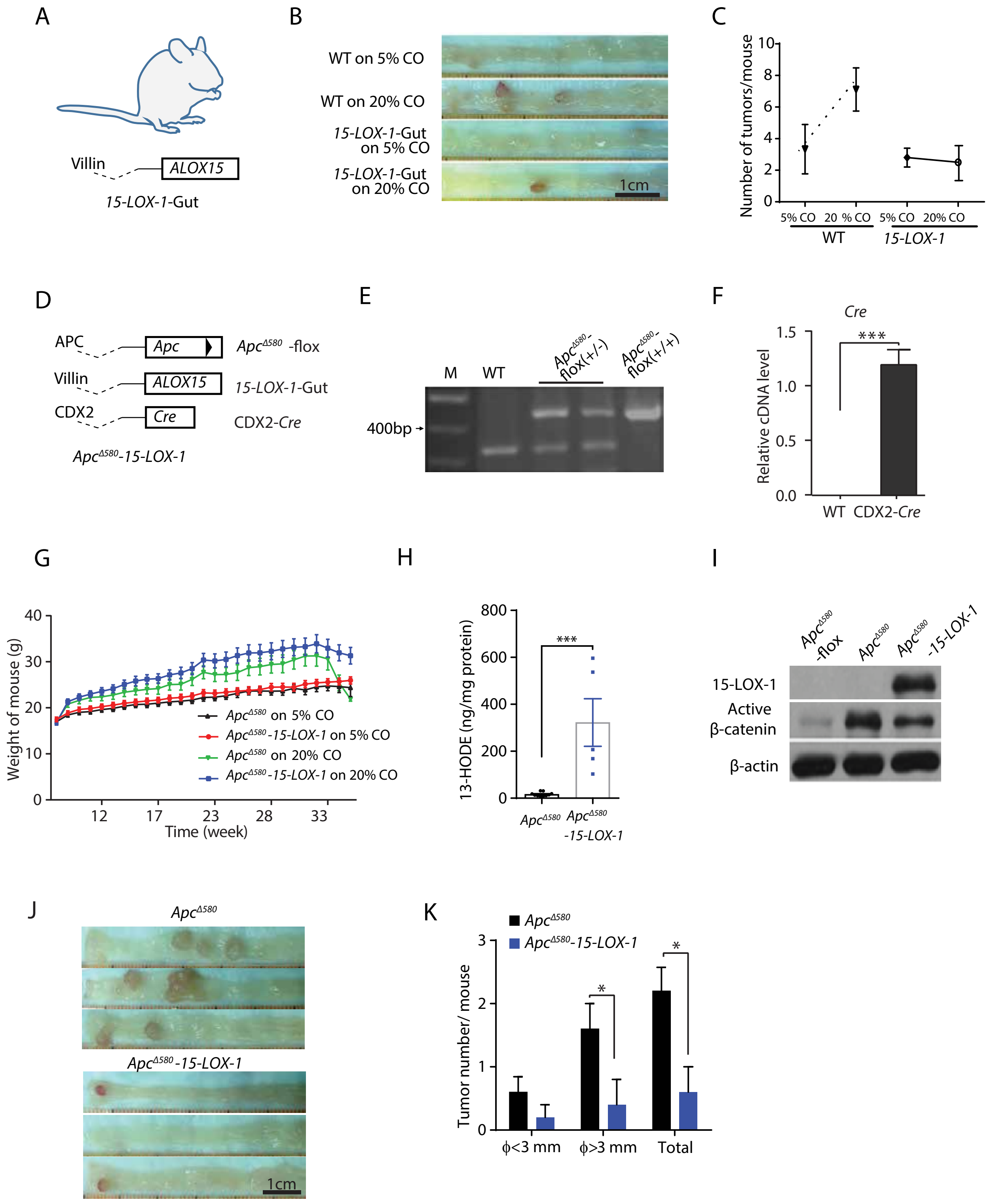

**Supplementary Figure 1. 15-LOX-1 suppresses linoleic acid (LA) promotion of AOM-induced colorectal cancer (CRC) and *Apc* mutation–induced CRC in mice.**

**(A)** Schematic diagram of villin-*15-LOX-1* (*15-LOX-1*-Gut) mice. Dashed lines indicate the promoter region, followed by the 15-LOX-1 gene coding region.

**(B, C**) *15-LOX-1*-Gut homozygotes and sex- and age-matched wild-type littermates (WT) were fed either 5% or 20% corn oil diet and then treated with 7.5 mg/kg AOM weekly via intraperitoneal injection for 6 consecutive weeks. Mice were killed 20 weeks after the last AOM injection and examined for tumor formation (n= 10 mice per group). Representative colonic images **(B)** and colonic tumor numbers per mouse **(C)** of the indicated mice groups.

**(D)** Schematic diagram of $Apc^{\Delta 580}$ *–15-LOX-1* mice. $Apc^{\Delta 580}$ mice were bred with *15-LOX-1*-Gut mice to produce $Apc^{\Delta 580}$*;15-LOX-1*-Gut, designated as $Apc^{\Delta 580}$*–15-LOX-1* mice.

**(E)** Genotyping of mice carrying $Apc^{\Delta 580}$-flox alleles, in which *Apc* exon 14 is flanked with *loxP* sites by regular polymerase chain reaction (PCR). *Apc* WT band: 320 bp. $Apc^{\Delta 580}$ flox band: 430 bp. M represents DNA ladder marker.

**(F)** *Cre* genotyping for CDX2-*Cre* mice. Genomic *Cre*-recombinase cDNA levels were measured by quantitative PCR. WT indicates no detectable genomic *Cre*-recombinase cDNA.

**(G)** $Apc^{\Delta 580}$ and $Apc^{\Delta 580}$ *–15-LOX-1* mice at 4 weeks were fed 5% or 20% corn oil diets, and the body weights of the mice were assessed every week until age 35 weeks. The body weight curves of the four mouse groups are shown (n=14-20 per group).

**(H, I)** Normal colonic epithelial cells of $Apc^{\Delta 580}$-flox, $Apc^{\Delta 580}$, and $Apc^{\Delta 580}$*–15-LOX-1* mouse groups were scraped and harvested for measuring 13-HODE levels by LC-MS/MS **(H)** and 15-LOX-1 and active β-catenin levels by Western blot **(I).**

**(J, K)** 15-LOX-1 expression inhibited CRC tumorigenesis in $Apc^{\Delta 580}$ mice. $Apc^{\Delta 580}$ mice and $Apc^{\Delta 580}$*–15-LOX-1* mice fed a standardized diet with fixed LA content (7% corn oil) were followed for 25 weeks and evaluated for tumor formation (n=5 per group). Representative colon photographs **(J)** and the colonic tumor numbers per mouse with tumor size distributions **(K)** of $Apc^{\Delta 580}$ and $Apc^{\Delta 580}$*–15-LOX-1* mice.

Scale bars: 1 cm (B and J). ∅: diameter.

Supplementary Figure 2

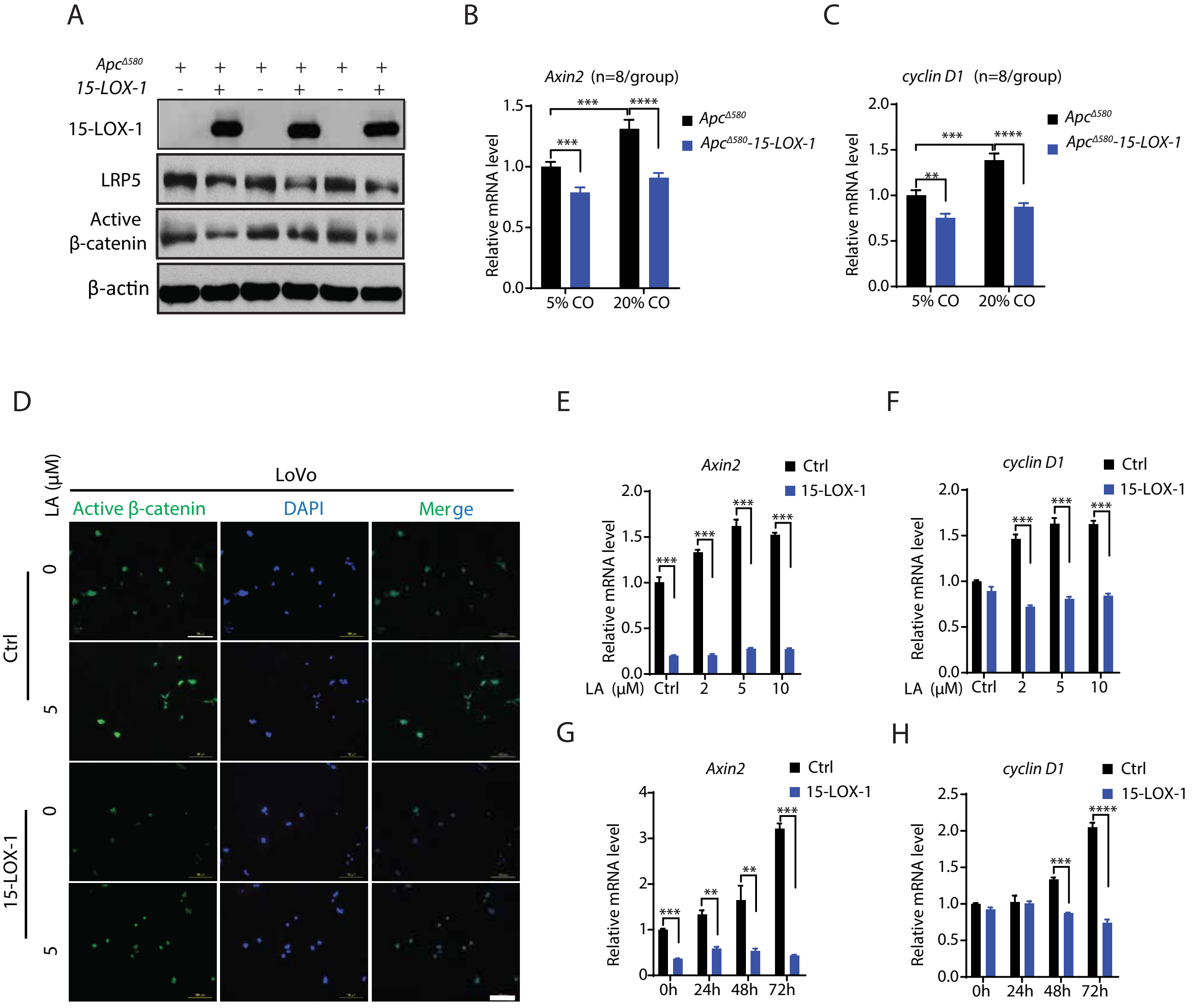

**Supplementary Figure 2. 15-LOX-1 suppresses LRP5 and active β-catenin expression in IECs of mice and human CRC cells.**

**(A)** LRP5 and active β-catenin protein levels in normal IECs of *Apc*$^{\Delta 580}$ and *Apc*$^{\Delta 580}$*-15-LOX-1* littermates at age 6-8 weeks were measured by Western blot.

**(B, C)** *Apc*$^{\Delta 580}$ and *Apc*$^{\Delta 580}$ *–15-LOX-1* mice at 4 weeks were fed 5% or 20% corn oil (CO) diets for 10 weeks. The mRNA expression levels of *Axin2* **(B)** and *cyclin D1* **(C)** in the IECs of the mice were measured by qRT-PCR.

**(D)** Immunofluorescence staining of active β-catenin in LoVo stably transduced with either control (Ctrl) or 15-LOX-1 lentivirus and treated with 5 μM LA for 48 hours. Representative immunofluorescence images are shown.

**(E-H)** *Axin2* and c*yclin D1* mRNA expression in LoVo cells stably transduced with either control (Ctrl) or 15-LOX-1 lentivirus and treated with LA at different concentrations in culture medium supplemented with 5% dialyzed FBS for 48 hours (**E,F**) or treated with 5 μM LA at different time points **(G, H),** measured by qRT-PCR.

Scale bars: 100 μm (F).

### Supplementary Figure 3

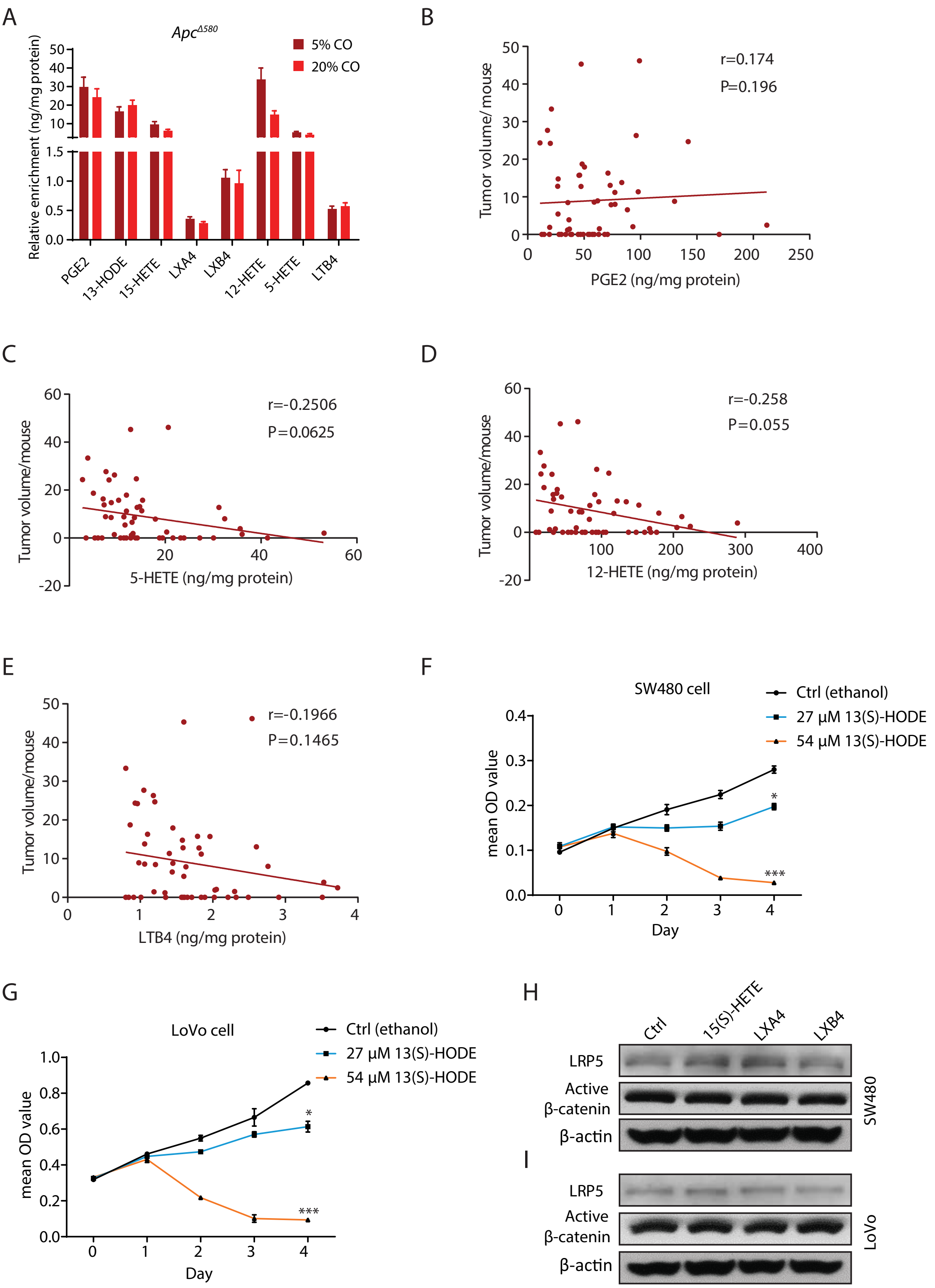

**Supplementary Figure 3. The profile analysis of 15-LOX-1–mediated eicosanoid metabolites in *Apc*$^{\Delta580}$ mice and 13(S)-HODE suppression of human CRC proliferation.**

**(A-E)** *Apc*$^{\Delta580}$ and *Apc*$^{\Delta580}$*-15-LOX-1* littermates at 4 weeks were fed either 5% or 20% corn oil diet for 10 weeks, killed at age 14 weeks, and examined for tumor formation (n=14 mice per group).

**(A)** Eicosanoid metabolite profile of the IECs of the indicated *Apc*$^{\Delta580}$ mouse groups were measured by liquid chromatography-tandem mass spectrometry (LC-MS/MS). **(B-E)** Spearman correlation analysis of colonic tumor volumes with levels of PGE2 **(B)**, 5-HETE **(C),** 12-HETE **(D**), or LTB4 **(E)** in IECs of all the examined *Apc*$^{\Delta580}$ and *Apc*$^{\Delta580}$*-15-LOX-1* mice (n=56).

**(F, G**) 13(S)-HODE suppresses human CRC proliferation. SW480 **(F)** and LoVo **(G)** cells were treated with 0, 27, and 54 µM 13(S)-HODE for 0, 1, 2, 3, and 4 days. The cell viability and proliferation were measured by MTT assay.

**(H, I)** SW480 **(H)** and LoVo **(I)** were treated with 20 µM 15(S)-HETE, 100 nM LXA4, and 200 nM LXB4 for 48 hours, and the cells were harvested and examined for LRP5 and active β-catenin expression by Western blot.

Supplement Figure 4

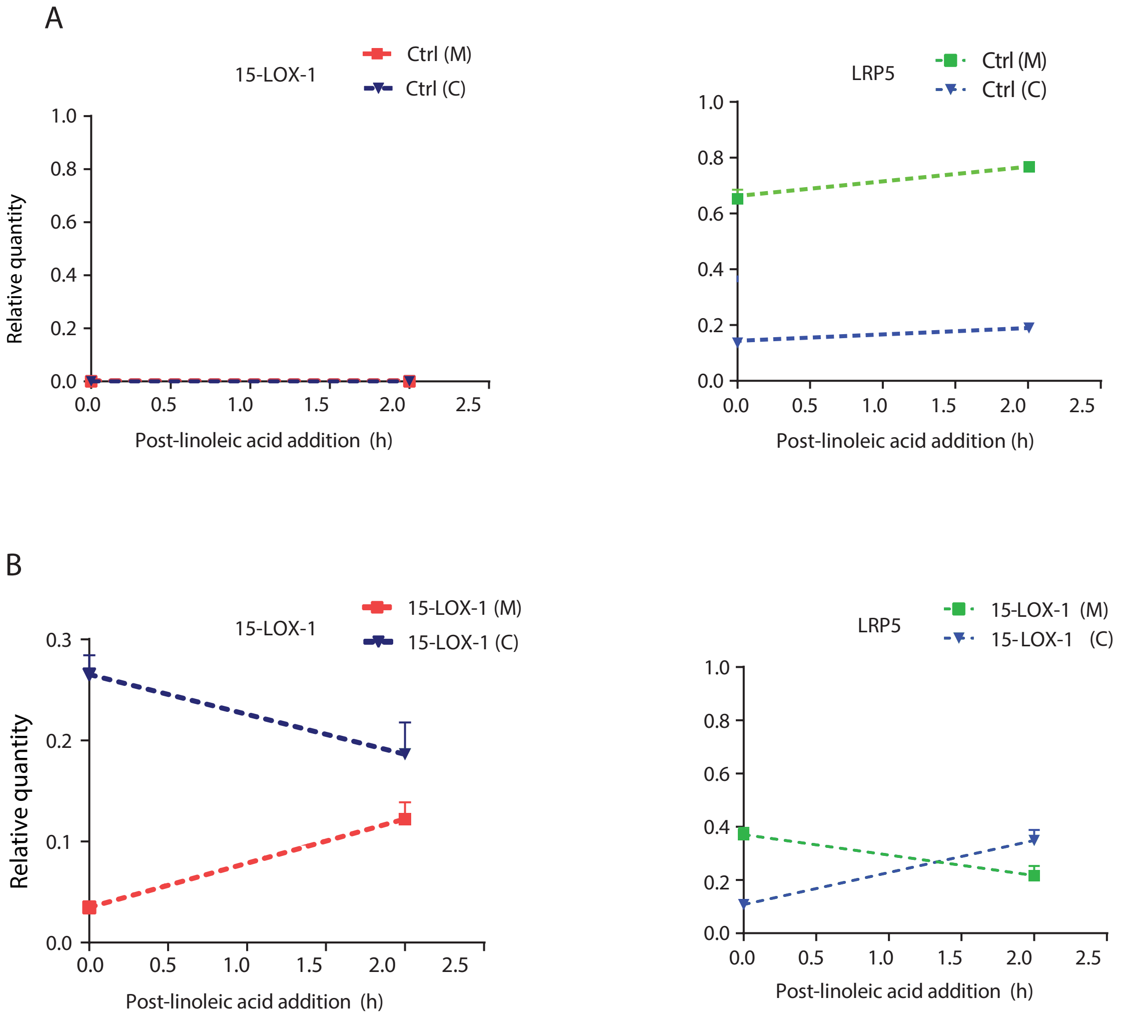

**Supplementary Figure 4. 15-LOX-1 decreases cell membranous LRP5 levels.**

(**A, B**) LoVo cells stably transduced with control (Ctrl) or 15-LOX-1 lentivirus treated with 10 μM LA for 2 hours were processed for cytoplasmic (C) and membranous (M) protein fractions, and then analyzed for LRP5 and 15-LOX-1 protein expression by Western blot. The ratios of protein band densities of cytoplasmic 15-LOX-1 or LRP5 over β-actin and membranous 15-LOX-1 or LRP5 over Na-K-ATPase in Ctrl **(A)** or 15-LOX-1 **(B)** lentivirus transduced LoVo cells corresponding to **Figure 5D**.

### Supplement Figure 5

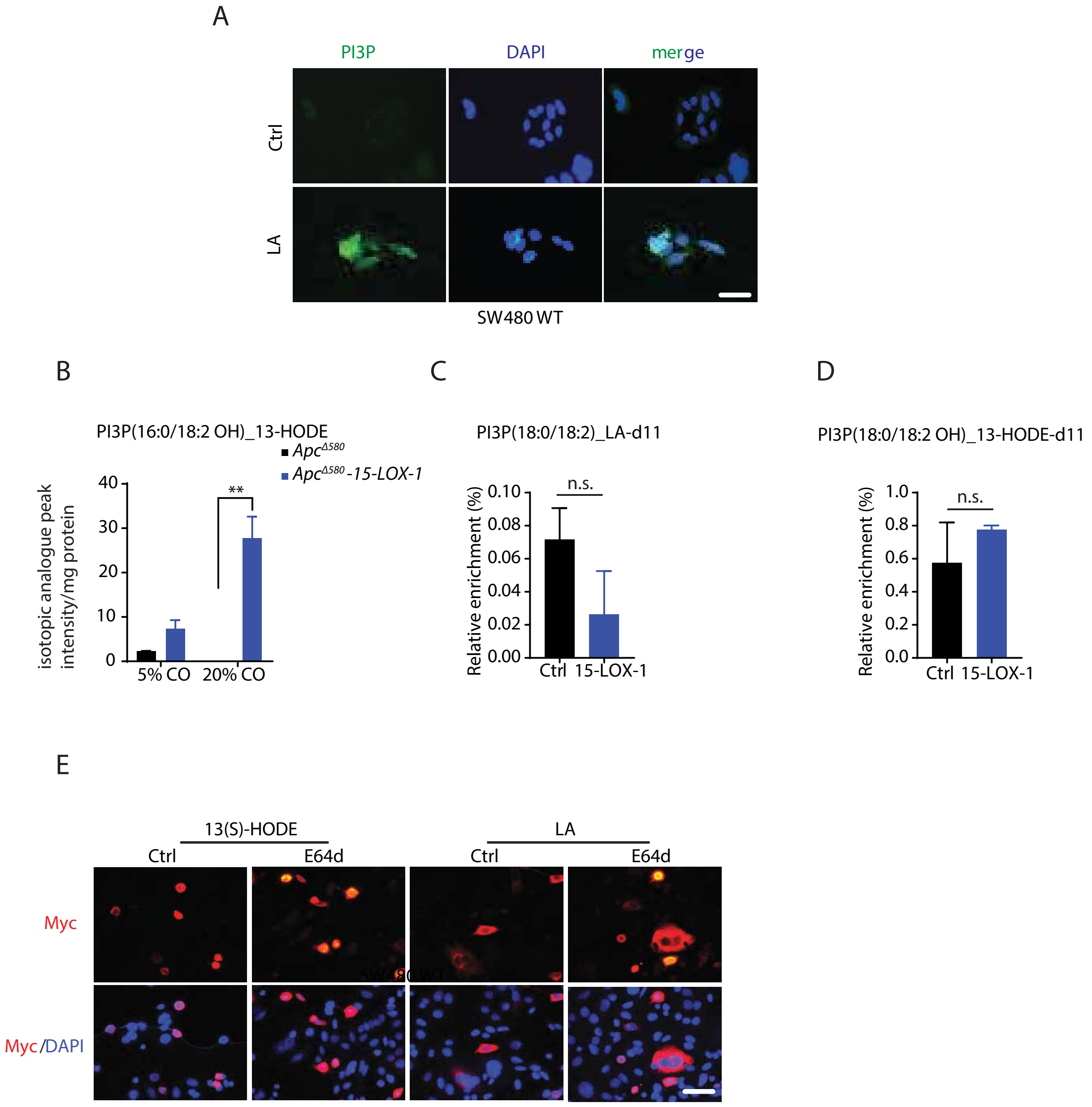

**Supplementary Figure 5. LA increases phosphatidylinositol 3-phosphate (PI3P) production and 15-LOX-1 increases PI3P_13-HODE production.**

**(A)** Immunofluorescence staining of PI3P in SW480 WT cells treated with 5 μM LA for 48 hours. Representative images are shown.

**(B)** *Apc*$^{\Delta 580}$ and *Apc*$^{\Delta 580}$*-15-LOX-1* littermates at age 4 weeks fed either 5% or 20% corn oil diet for 10 weeks were killed at age 14 weeks. The IECs of these mice were scraped and examined for patterns of PI3P incorporated with (16:0/18:2 OH)_13-HODE by LC-HRMS (n=3-4 mice per group).

**(C, D)** SW480 cells stably transduced with control (Ctrl) or 15-LOX-1 lentivirus were treated with 100 μM LA-d11 for 36 hours in culture medium containing 5% dialyzed FBS. PI3P profile was traced with d11-labeled LA by LC-HRMS. The percentage of d11-labeled PI3P incorporated with (18:0/18:2)_LA over PI3P incorporated with total (18:0/18:2)_LA **(C)**, or d11-labeled PI3P incorporated with (18:0/18:2 OH)_13-HODE over PI3P incorporated with total (18:0/18:2 OH)_13-HODE **(D),** was calculated and presented.

**(E)** SW480 cells transfected with Myc-tagged LRP5 expression vector for 48 hours, then treated with E-64d or solvent for 1 hour, followed by treatment of LA (100 μM) or 13(S)-HODE (27 μM) for an additional 4 hours. The cells were harvested and processed for immunofluorescence staining using anti–c-Myc antibody. Representative images are shown.

n.s. means no significance difference. Scale bars: 100 μm (A and E).
